# ALS molecular subtypes are a combination of cellular, genetic, and pathological features learned by deep multiomics classifiers

**DOI:** 10.1101/2024.07.19.603731

**Authors:** Kathryn O’Neill, Regina Shaw, Isobel Bolger, NYGC ALS Consortium, Oliver Tam, Hemali Phatnani, Molly Gale Hammell

## Abstract

Amyotrophic Lateral Sclerosis (ALS) is a complex syndrome with multiple genetic causes and wide variation in disease presentation. Despite this general heterogeneity, several common factors have been identified. For example, nearly all patients show pathological accumulations of phosphorylated TDP-43 protein in affected regions of the motor cortex and spinal cord. Moreover, large patient cohort studies have revealed that most patient samples can be grouped into a small number of ALS subtypes, as defined by their transcriptomic profiles. These ALS molecular subtypes can be grouped by whether postmortem motor cortex samples display signatures of: mitochondrial dysfunction and oxidative stress (ALS-Ox), microglial activation and neuroinflammation (ALS-Glia), or dense TDP-43 pathology and associated transposable element de-silencing (ALS-TE). In this study, we have built a deep layer ALS neural network classifier (DANcer) that has learned to accurately assign patient samples to these ALS subtypes, and which can be run on either bulk or single-cell datasets. Upon applying this classifier to an expanded ALS patient cohort from the NYGC ALS Consortium, we show that ALS Molecular Subtypes are robust across clinical centers, with no new subtypes appearing in a cohort that has quadrupled in size. Signatures from two of these molecular subtypes strongly correlate with disease duration: ALS-TE signatures in cortex and ALS-Glia signatures in spinal cord, revealing molecular correlates of clinical features. Finally, we use single nucleus RNA sequencing to reveal the cell type-specific contributions to ALS subtype, as determined by our single-cell classifier (scDANCer). Single-cell transcriptomes reveal that ALS molecular subtypes are recapitulated in neurons and glia, with both ALS-wide shared alterations in each cell type as well as ALS subtype-specific alterations. In summary, ALS molecular subtypes: (1) are robust across large cohorts of sporadic and familial ALS patient samples, (2) represent a combination of cellular, genetic, and pathological features, and (3) correlate with clinical features of ALS.

Figure 0:
Graphical Abstract - ALS molecular subtypes are a combination of cellular, genetic, and pathological features learned by deep multiomics classifiers.

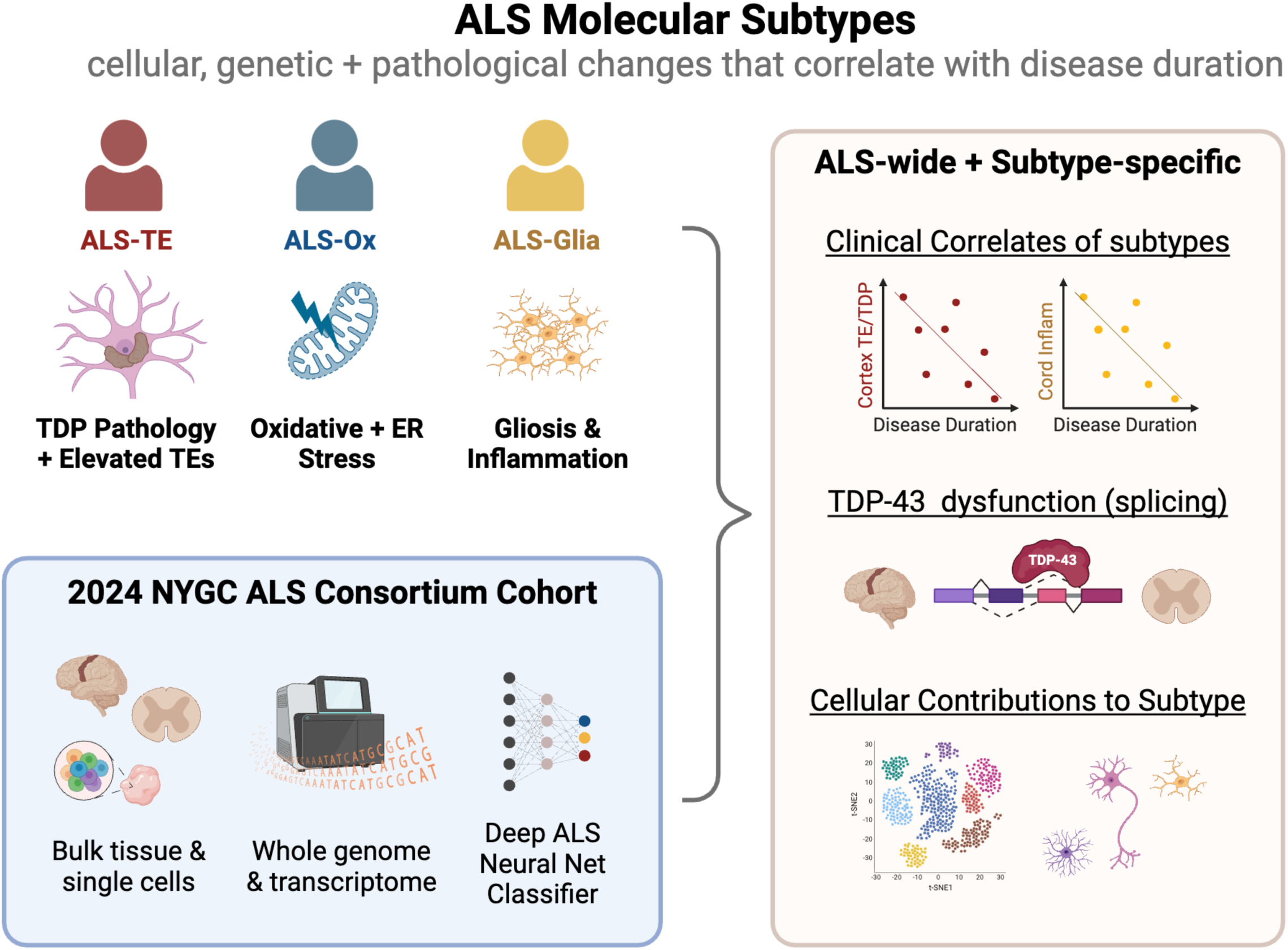

## Introduction

Amyotrophic Lateral Sclerosis (ALS) is a fatal neurodegenerative disease characterized by the progressive loss of upper and lower motor neurons of the motor cortex and spinal cord^1^. These motor neurons control voluntary motor function, which typically leads to total paralysis and eventual death within 2-5 years of diagnosis^2^. Across people living with ALS (pALS), there is a high degree of variability in terms of the site of disease onset, the age at diagnosis, the rate of disease progression, and the relative involvement of upper and lower motor neuron associated symptoms. Despite this heterogeneity, nearly all ALS patients show pathological accumulations of phosphorylated TDP-43 protein in spinal cord and motor cortex tissues^3,4^. TDP-43 pathology is present in most cases with known ALS causal mutations, typically called “familial ALS”^5^. At least 40 and as many as 90 genetic mutations have been associated with ALS^6^. These ALS associated genes can generally be grouped by their inferred roles^5^ in RNA processing, protein homeostasis, neuroinflammation, and cytoskeletal dynamics. TDP-43 pathology is additionally present in nearly all pALS cases without a family history of disease or known causal mutation, often called “sporadic ALS”, with approximately 90% of all ALS cases estimated to have the sporadic form^7^.

Several other neurodegenerative diseases also present with pathological inclusions of TDP-43^8–10^, most prominently in the TDP-43 subset of patients with frontotemporal dementia (FTD-TDP)^10^. ALS and FTD-TDP share many common genetic factors and a high degree of co-morbidity, such that these two diseases are understood to exist on a spectrum^11^. For example, patients with inherited mutations in the C9orf72 gene may present with pure ALS, pure FTD, or a combination of both^12^. In addition to this shared pathology, other molecular alterations are commonly seen across tissues from patients along the ALS/FTD spectrum. Previous work from our own laboratories using genomics profiling of a large ALS patient cohort, the NYGC ALS Consortium, revealed that most ALS patient tissues can be grouped into a small number of ALS subtypes, as defined by their transcriptomic profiles^1^. These ALS molecular subtypes can be grouped by whether *post mortem* cortex samples display signatures of: mitochondrial dysfunction and oxidative stress (ALS-Ox), microglial activation and neuroinflammation (ALS-Glia), or dense TDP-43 pathology and associated transposable element de-silencing (ALS-TE). These molecular subtypes identified from cortical transcriptome data were validated at the protein level via immunohistochemistry (IHC) stains for proteins reflective of that ALS subtype. For example, ALS tissue samples with an ALS-Glia expression profile were marked by increased density of hypertrophic, IBA1 positive, pro-inflammatory microglia. In contrast, pALS cases with an ALS-TE subtype instead showed dense clusters of cells with pTDP-43 positive aggregates present in cortical tissues. Work from other groups on independent ALS patient cohorts has since replicated these findings in motor cortex and detected molecular subtypes in regions the central nervous system.^13^ Finally, work from other groups on the same cortical samples as those originally presented by the NYGC ALS Consortium cohort also replicated the original findings for ALS molecular subtypes^14^. Taken together, these studies have confirmed the presence of ALS molecular subtypes in frontal and motor cortex^1,13,14^, shown that these subtypes may be detectable in peripheral biofluids^13^, and detected an association between certain ALS subtypes and clinical correlates^13,14^ such as age of onset and disease duration.

Previous work on ALS molecular subtypes has left open several questions about the nature and implications of each subtype. First, subtypes have so far been identified only in bulk RNA-seq transcriptomics studies, abrogating our ability to determine cell-type-specific gene expression alterations. Moreover, changes in bulk expression patterns might be explained more simply by alterations in cell type composition of the tissue. Second, these studies profiled only a few hundred ALS patients in each cohort, and would have been underpowered to detect rare subtypes. Lastly, these studies have not yet integrated spinal cord tissues with the cortical expression patterns, despite the importance of spinal motor neuron degeneration in ALS. In this study, we have addressed these gaps in our understanding of ALS molecular subtypes. We have integrated single nucleus transcriptome profiles to determine the cell type contributions to ALS molecular subtypes and show that ALS subtypes cannot be explained purely by changes in cell composition. Using an updated set of NYGC ALS Consortium samples that more than triples the size of the original study, we show that ALS molecular subtypes are robust across a large patient population from additional clinical centers and found no evidence of new subtypes in this greatly expanded study. We have additionally integrated matched cortical and spinal cord tissues, outlining the extent to which subtypes are present across these very different tissue types. To support these findings, we present a deep ALS neural net classifier (DANCer) that is able to take in newly sequenced profiles and assign an ALS molecular subtype with high accuracy and precision. DANCer can additionally work on single-cell/nucleus transcriptomes in its single-cell mode (scDANCer), as validated using samples with matched bulk and single-nucleus transcriptome profiles.

## Results

### 1. Expansion of the NYGC ALS Consortium Patient Samples

The NYGC ALS Consortium study has greatly expanded since 2019, adding new whole genome and transcriptome samples from several CNS tissues from additional pALS samples and from neurological and non-neurological control subjects. Since the original ALS molecular subtype study, the NYGC ALS consortium has more than tripled the number of available samples to provide whole genomes and transcriptomes for 719 cortical samples from 398 pALS and control individuals (286 from pALS, 53 neurological controls, and 59 non-neurological controls), see Figure 1A. The ALS Consortium samples were collected from 10 clinical centers and partnering institutions. The patients who donated these samples are relatively balanced in terms of patient sex (Table S1), with a median age at diagnosis of 64 (Table S1), a median disease duration of 32 months post-diagnosis (Table S1), and were derived from a largely sporadic cohort with few patients carrying known mutations of large effect (Fig 1B). Most of the ALS/FTD samples in the NYGC ALS Consortium cohort were obtained from patients (n = 286) with a diagnosis of ALS; 8% were also diagnosed with the related disorder frontotemporal dementia (FTD), see Figure 1A. Among the ALS Spectrum cases (ALS with or without FTD), 14% of the NYGC consortium cohort carried repeat expansion mutations in the C9orf72 gene (Fig 1B), 2% carried mutations in SOD1, and 1% of ALS cases carried mutations in the FUS, ATXN2, or OPTN genes. All other mutations were present in a single case and involved mutations in the genes: NEFH, VCP, PNPLA6, FIG4, UBQLN2, and PFN1. The distribution of pALS donors across site of onset, age of onset, and disease duration are similar to that seen in previous releases from the NYGC ALS Consortium^15–17^ and to larger estimates of disease presentation in additional ALS consortium cohorts, such as CREATE^18^, Project Mine^19^, Answer ALS^20^.

**Figure 1:**
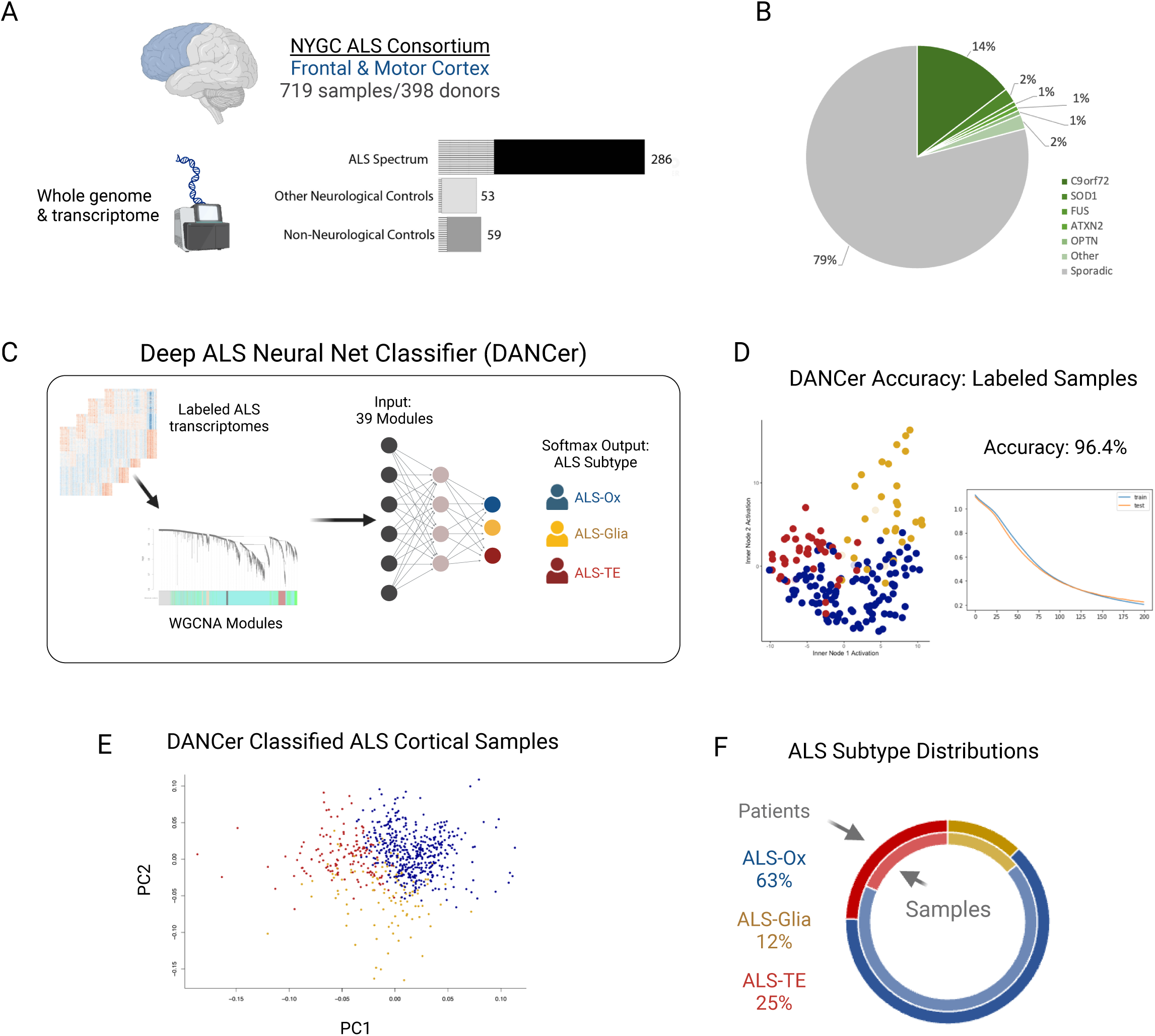
A Deep ALS neural net classifier (DANCer) accurately assigns ALS molecular subtypes to an expanded NYGC ALS Consortium cohort of ALS cortical samples. (A) The 2023 NYGC ALS Consortium genomics profiling cohort consists of 719 samples of frontal and motor cortex from 398 donors. Whole genome and transcriptome libraries were generated for 286 donors with ALS/FTD Spectrum disorders, 53 Non-ALS Neurological controls, and 59 Non-Neurological controls. (B) For cases on the ALS Spectrum, 79% of all tissues were from sporadic ALS donors while 15% carried mutations in C9orf72, 4% carried mutations in SOD1, and only 3 other gene mutations were present in more than one case (FUS, ATXN2, and OPTN). (C) A Deep ALS Neural Net Classifier (DANCER) was developed to learn ALS subtype assignments using previously published datasets^1^ as the training set. DANCer takes whole transcriptome data as input, converts those gene count tables into eigengenes, and assigns ALS subtypes using a multilayer, fully connect, neural network. (D) DANCer accuracy was assessed using 20-fold cross-validation on the labeled data^1^, reaching 96.4% after 200 training epochs. Accuracy is graphically displayed for ALS-Ox (blue), ALS-Glia (gold), and ALS-TE (red) samples in the latent variable projection space, where solid circles represent accurately called subtypes and transparent circles represent incorrect calls. (E) DANCer classification was then applied to the 719 samples in the NYGC ALS Consortium cohort, with a graphical display of sample distributions in PCA space. (F) The distribution of ALS cases across the 3 ALS subtypes (63% ALS-Ox, 12% ALS-Glia, 25% ALS-TE) is similar to what has previously been seen in smaller ALS cohort^1,13,14^.

### 2. A Deep ALS Neural Net Classifier (DANCer) Accurately Assigns ALS Molecular Subtypes

The original identification of ALS Molecular Subtypes was enabled by a de novo clustering and pattern discovery method called non-negative matrix factorization (NMF)^21^. While NMF is a powerful technique to define new molecular pathways associated with samples in bulk transcriptomic data, it is impractical to apply in the context of continuously expanding collections of data. As NMF is a factorization method, de novo pattern discovery within the entire sample cohort must be re-calculated each time new data is added. To address these limitations, we developed a classifier using an artificial neural network architecture to accurately predict ALS molecular subtype assignment in new incoming samples. Here we present our Deep ALS Neural net Classifer (DANCer).

Initially, we sought to determine how easily the transcriptomic structure of molecular subtype could be captured by neural networks (NNs). To do this we constructed an autoencoder which accepted the same input transcriptomes as those originally used for NMF discovery in the Tam et al dataset^1^ – the top 5000 genes ranked by median absolute deviation (MAD5000). The architecture consisted of 7 layers of 5000, 256, 256, 2, 256, 256, and 5000 nodes respectively, as seen in Figure (S1A). The autoencoder, and all subsequent NN architectures, was built using the Keras library^22^, a loss function that calculated mean squared error (MSE) between input and output, random dropout, and early stopping to prevent overfitting. We monitored for convergence within 200 training epochs with early stopping by reduced loss according to MSE. We then visualized the firing rates on the inner bottleneck nodes for each ALS transcriptome sample in the fully trained model, in order to determine what the autoencoder model was learning. Figure S1A shows these bottleneck node firing weights colored by the previously labeled ALS molecular subtype assignments^1^. In Figure S1A, ALS molecular subtypes clearly occupied disparate locations in this compressed two-dimensional representation, indicating subtype structure was readily captured by neural networks.

Convinced that ALS Molecular Subtypes were learnable by neural network (NN) architectures and reproducible across machine learning methods (NMF and NNs), we then sought to convert the autoencoder into a fully trained classifier (Fig 1C). To reduce noise and redundancy within the transcriptome datasets, we first used weighted gene correlation network analysis (WGCNA)^23^ as a dimensionality reduction method that enabled us to efficiently represent the information from all expressed genes in our training dataset. Briefly, WGCNA clusters a gene set into modules of co-expressed genes, or groups of genes with similar expression profiles as determined by a distance metric. Each of these modules are represented by a single value named an eigengene, or the first principal component of all the genes in a module. One module/eigengene was excluded as it consisted entirely of sex-biased genes. The remaining 39 eigengenes were used as inputs for a multilayer feedforward perceptron NN (Fig 1C), which was trained to reproduce the labeled ALS molecular subtypes outlined in our original study^1^. Following 200 training epochs, with 20-fold cross validation, the classifier reached 96.4% classification accuracy on the original labeled datasets (figure 1D). The accurately assigned ALS samples are represented by solid dots in the autoencoder latent space scatterplot (Fig 1D), while the small number of incorrectly assigned samples are represented by translucent dots. Additional details about the training protocols, parameters used in the models, and generation of an autoencoder latent space, can be found in the methods section and supplementary Figure S1. The code for DANCer is available at Github at the following link https://github.com/mhammell-laboratory/DANCer.

### 3. DANCer Classifications show ALS molecular subtypes are robust across an expanded ALS cohort

We next applied the DANCer classification algorithm to the updated 2023 release of frontal and motor cortex samples from the NYGC ALS Consortium. The input data for DANCer consisted of whole transcriptome RNA-seq datasets from 719 frontal and motor cortex samples of 398 subjects, as described above, and are visualized in a PCA representation color-coded by ALS subtype (Fig 1E). The distribution of samples into ALS molecular subtypes was broadly similar to that seen in the previous NYGC ALS Consortium release^1^, with 63% of pALS donors in the ALS-Ox group, 12% of pALS donors in the ALS-Glia group, and 25% classified as ALS-TE (Fig 1F).

These sample level assignments were then converted to a patient-level assignment for donors with more than one sample, using a majority call for any patients with discordant calls between samples as previously described^1^. The updated set of NYGC ALS Consortium pALS patient subtypes were classified as 17% ALS-TE, 66% ALS-Ox, and 16% ALS-Glia (Fig 1F). This distribution is broadly similar to that seen previously when the ALS subtypes were originally described, with only small changes in the number of ALS/FTD subjects with the inflammatory ALS-Glia subtype (16% as compared to 19%) and the ALS-TE subtype that shows markers of TDP-43 dysfunction and transposable element up-regulation (17% as compared to 20%). These small changes may reflect differences in patient populations sampled and inclusion of more samples from patients with co-morbid FTD. When looking just at patients with a pure FTD diagnosis, out of all those on the ALS/FTD spectrum, the distribution shifts more towards the ALS-TE & ALS-Glia subtypes with 43% of the FTD patients assigned as ALS-TE, 27% as ALS-Ox, and 30% as ALS-Glia (Table S1C).

Using only ALS Spectrum patients, we next explored whether any biological and/or technical co-variates might play a role in sample subtype assignments. We found no significant difference between ALS-Glia or ALS-TE subtypes and controls for technical variables such as sample RNA integrity values (RIN) values or tissue pH. The ALS-Ox subtype samples showed slightly higher RIN values when compared to controls: ALS-Ox mean RIN = 7.1, control mean RIN = 6.55, Wilcoxon P<6.5e-7 (see Table S2a). We also explored biological co-variates, such as: sex, site of onset, ALS/FTD diagnosis, age at diagnosis, and known ALS mutations. We found no association for most of these biological variables (Supp Table 2b), though we acknowledge that we are under-powered to assess associations with mutational status. We note that the ALS-Ox subtype was significantly enriched for patients with a pure ALS diagnosis (odds ratio 3.2, Fisher’s P<2.3e-6) and depleted for patients with a pure FTD diagnosis (odds ratio 0.2, Fisher’s P<9.1e-4). In contrast, the ALS-TE subtype was enriched among FTD patients (odds ratio 2.6, Fisher’s P<0.065) and depleted for patients with a pure ALS diagnosis (odds ratio 0.28, Fisher’s P<3.4e-6).

Finally, to ensure that no new subtypes are present in this larger cohort of NYGC ALS Consortium patients, we re-ran the de novo pattern finding algorithms that were originally used to define ALS subtypes, non-negative matrix factorization (NMF)^21^. NMF again returned an optimal clustering of k=3, and the three clusters returned showed a similar set of class assignments as previously described (see Supp Fig S2). Taken together, these results suggest that ALS subtypes are robust in terms of distributions and defining molecular characteristics across large patient cohorts originating from many different geographical areas and clinical centers. This release represents the largest ALS multi-omics data cohort to date and allows for robust determination of the correlation between molecular genomic profiles and clinical metadata.

### 3. ALS Spinal Cord samples recapitulate the ALS-Ox and ALS-Glia subtypes

The NYGC ALS Consortium sample set also contains transcriptomes for 664 ALS and control patient samples from cervical, thoracic, and lumbar regions of the spinal cord. Of these, 553 were available from subjects for which we also have cortical tissue profiles. To determine whether the ALS subtypes would also be present in spinal cord samples, we ran the NMF matrix factorization algorithms on the full set of ALS and control spinal cord samples. NMF returned an optimal clustering of k=2, with just two dominant transcriptional patterns present in spinal cord samples (Fig 2A and Supp Fig S3). When we explored the genes that distinguished these two transcriptional clusters in spinal cord, a clear pattern emerged (Fig 2B): oxidative stress genes were elevated in the first group (NMF1) and inflammatory pathways in the second group (NMF2). Specifically, gene set enrichment analysis (GSEA)^24^ returned pathways such as Complement Cascades, Inflammatory Response, and Interferon Gamma Response as marking NMF2 (Fig 2B, gold) while Neuroactive Ligands and Receptors, Oxidative Phosphorylation, and Parkinson’s Disease genes marked the NMF1 spinal group (Fig 2B, blue). Furthermore, specific markers of pro-inflammatory astrocytes and microglia were significantly enriched in the NMF2/inflammatory spinal cord samples (Fig 2C), including those describing: Disease Associated Astrocytes^25^, Astrocytes induced via LPS or oxidative stress^26^, and pro-inflammatory microglia associated with hyper-active TREM2 signaling^27^. The marker genes identified from microglia isolated from SOD1 mutant mice^28^ showed the strongest enrichment score (NES=2.45) for ALS patient spinal cord samples in the inflammatory group, though we note that very few of these patients carried SOD1 mutations. To better reflect the genes expressed by these two spinal cord transcriptome groups, we re-named the first (NMF1/blue) group “ALS-Ox-Cord” and the second (NMF2/gold) group “ALS-Glia-Cord” (Fig 2A).

**Figure 2:**
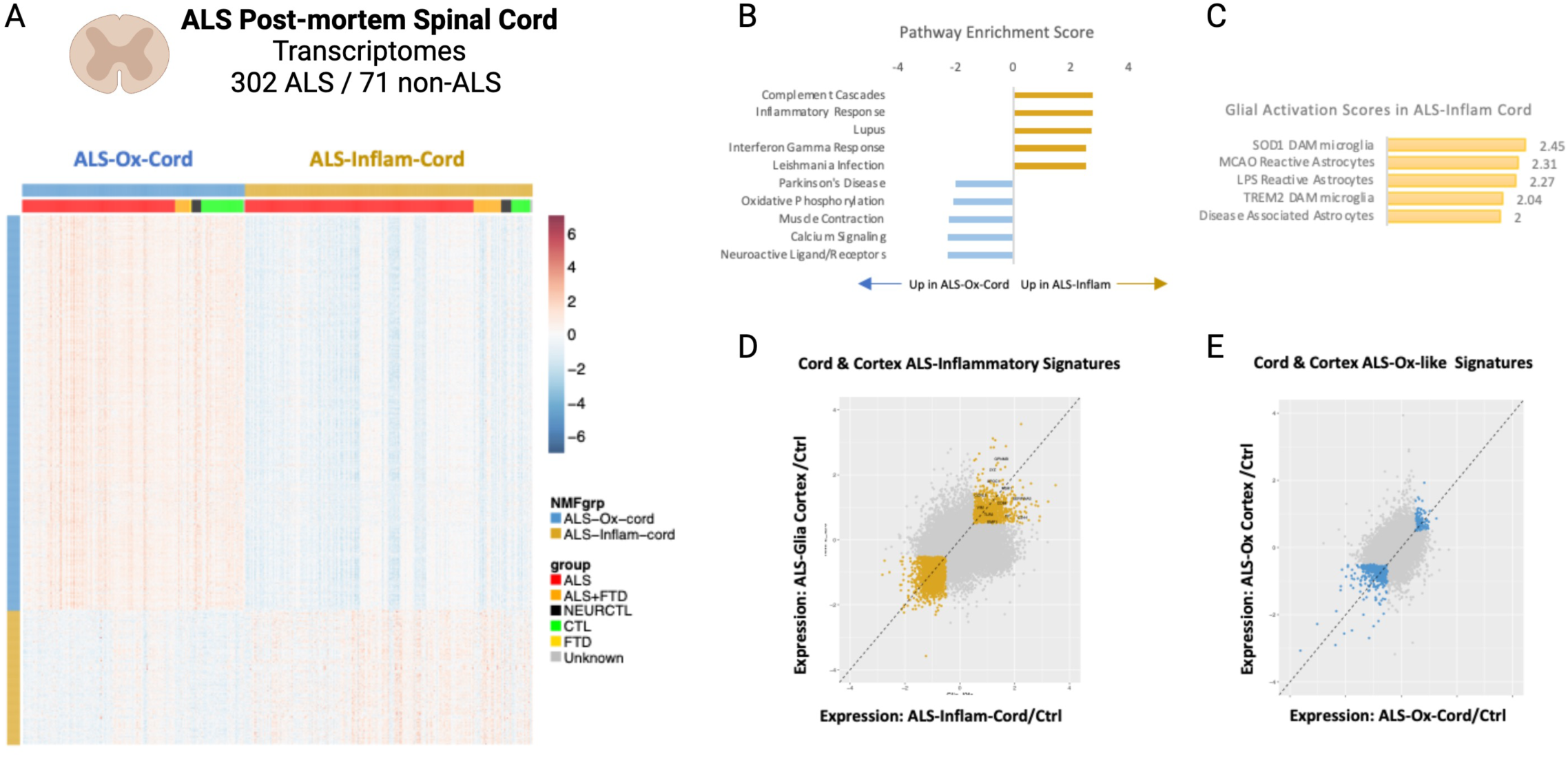
ALS Spinal Cord transcriptional patterns are dominated by inflammatory & oxidative stress pathways, showing strong overlap with ALS-Glia and ALS-Ox cortical subtypes. (A) The 2023 NYGC ALS Consortium cohort additionally contains 302 ALS and 71 non-ALS spinal cord samples. Non-negative Matrix Factorization (NMF) was applied to these samples to ascertain whether molecular subtypes would emerge in these samples as well. NMF returned k=2 transcriptome subtypes, with marker genes that indicated either oxidative stress (blue) or inflammatory (gold) gene expression patterns. (B) Gene Set Enrichment Analysis (GSEA) of these NMF marker genes returned enrichments for pro-inflammatory pathways for the ALS-inflammatory-cord samples (gold), while ALS-Ox-cord samples were enriched for patterns of oxidative phosphorylation and Parkinson’s Disease (blue). (C) GSEA was also run on glial activation markers from several published studies, with the greatest enrichment seen for markers of SOD1 mutation induced disease associated microglia^28^ (SOD1-DAM). (D) To stress the similarity of the inflammatory signatures in spinal cord and cortex samples, we’ve plotted the fold changes for all genes in ALS-Glia cortical samples vs controls (y-axis) against all measured fold changes for ALS-Glia-cord samples vs. controls (x-axis). (E) To stress the similarity of the oxidative stress signatures in spinal cord and cortex samples, we’ve plotted the fold changes for all genes in ALS-Ox cortical samples vs controls (y-axis) against all measured fold changes for ALS-Ox-cord samples vs. controls (x-axis).

Given the similarities in gene pathways identified for the two ALS spinal cord groups (oxidative stress and inflammatory pathways) and two of the subtypes previously identified from ALS cortex samples (ALS-Ox and ALS-Glia), we next sought to understand whether the spinal cord and cortical samples are recapitulating the same ALS associated molecular subtypes as previously identified in cortex. We first looked at the inflammatory signatures expressed in the ALS-Glia-Cord (gold) samples. We compared the average fold change for all genes in ALS-Glia-cord samples from ALS patient spinal cord samples to the average fold change for all genes in ALS-Glia cortex samples, relative to controls (Fig 2D). We find a significant correlation between the expression of these genes in ALS patient cortex and spinal cord tissues (r=0.38, P<2.2E-16) despite the large differences in cellular composition between these two tissues. We repeated the same comparisons for ALS-Ox signatures from the cortex samples and the ALS-Ox-like signatures from the spinal cord (Fig 2F). Again, we see a significant correlation for altered gene expression in these two tissues (r=0.49, P<2.2E-16). This suggests that the ALS-Glia and ALS-Ox cortical subtype patterns are recapitulated in the spinal cord and conserved throughout the motor system.

Previous reports of inflammatory signatures that mark ALS patient samples identified several genes that were dysregulated at both the transcriptional and protein level in ALS spinal cord and in samples from ALS cerebrospinal fluid (CSF)^16^. We can recapitulate these findings in the ALS-Inflammatory group and find that these genes are also amongst the most strongly correlated between their elevation in the ALS-Glia group in the spinal cord and the ALS-Glia group in the cortex. Several of these genes are marked in Figure 2D, including GPNMB, SERPINA3, and CHIT1 and listed in Supplementary Table S4G. The fact that inflammatory signatures are more prevalent in spinal cord over ALS-Ox-like signatures likely explains why that same signature would be recapitulated when comparing all ALS samples to controls, as was done by Humphrey et al^16^. However, the ALS-Ox-cord samples are not comparable to controls and instead show thousands of differentially expressed genes when compared to controls (Supp Table S4I).

### 4. Integrative analysis across spinal cord & cortex reveals correlates of ALS molecular subtypes with TDP-43 dysfunction

Given the similarities between ALS-Ox and ALS-Glia signatures in spinal cord and cortex, we next explored whether each patient would show a single concordant pattern in both cord and cortical tissues (Fig 3A and Table S5). There were 151 donors with both cortex and spinal cord tissues profiled. A majority of the patients with the ALS-Glia/inflammatory signature in the cortex also showed ALS-Glia/inflammatory signatures in the spinal cord (16/20=80%). However, nearly all ALS donors showed evidence of inflammatory pathway signatures in spinal cord tissues (103/151=68%). This point emphasizes the stark differences in the relative frequency of inflammatory signatures in the cortex and spinal cord. As noted above, ALS-Glia subtype patients formed the smallest group of all subtypes in the NYGC ALS Consortium Cohort in cortex (12%), yet formed the majority of spinal cord subtypes (68%).

**Figure 3:**
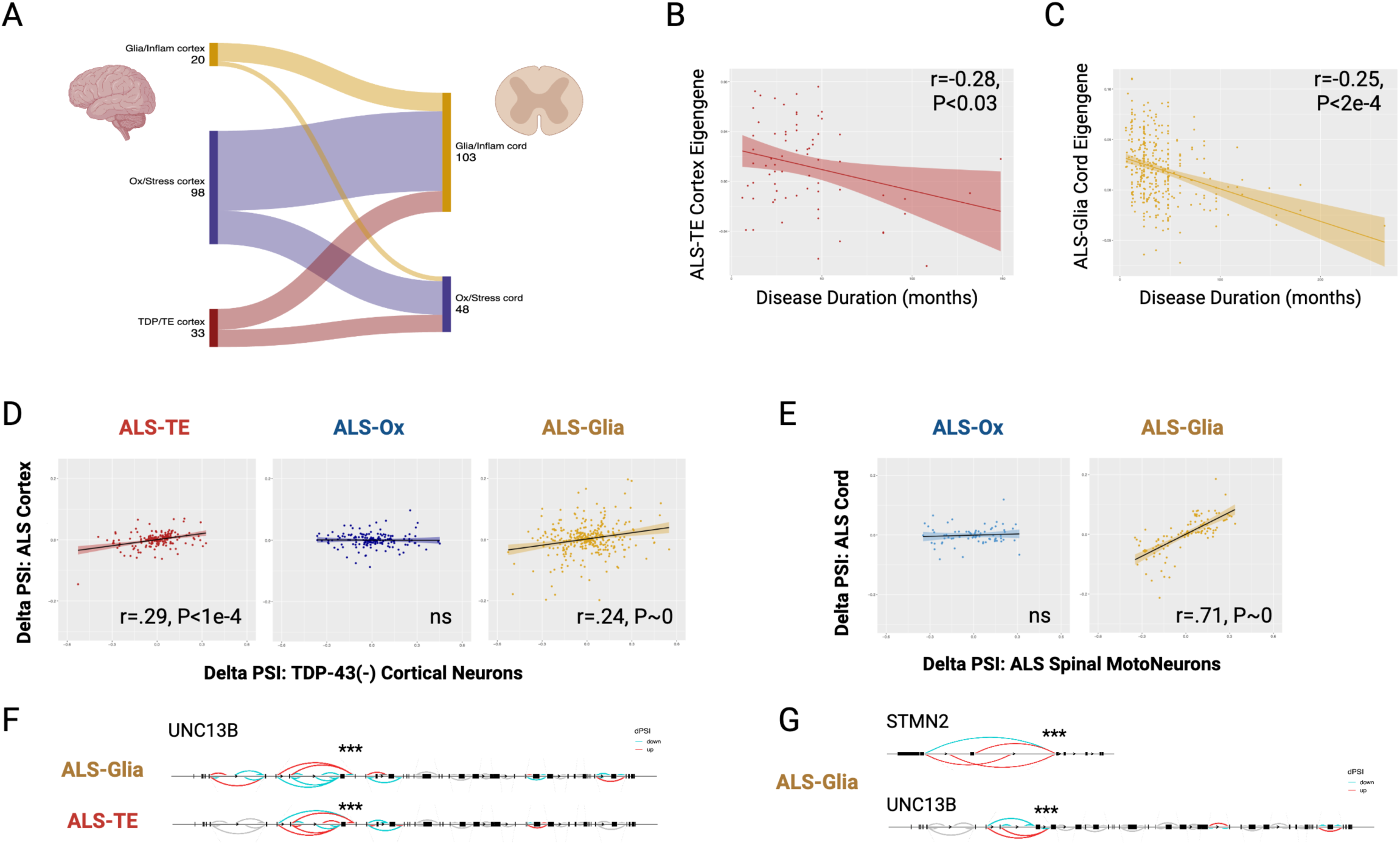
Integrative analysis across cortical and spinal cord tissues shows correlates of ALS molecular subtypes with Clinical Features. (A) For 200 of the ALS cases in the NYGC ALS Consortium cohort, we have matched spinal cord and cortex samples. This enabled us to look at the relative distribution of ALS-Ox, ALS-Glia, and ALS-TE samples in each region. We see differing patterns, where the ALS-Ox subtype dominated the calls for the cortex (139/200= 70%), but were present in a minority of the spinal cord cases (74/200=37%). Conversely the ALS-Glia subtype dominates in the spinal cord (126/200=63%), but was the smallest subset of cortex. The ALS-TE subtype is only detectable in cortex. (B) The ALS-TE subtype shows a strong negative correlation between ALS-TE gene expression signatures and disease duration, indicating shorter disease duration for these ALS cases. (C) Inflammatory gene expression signatures in the spinal cord were significantly negatively correlated with disease duration. (D) The ALS-TE subtype showed the strongest signature for TDP-43 dysfunction, as measured by comparing splicing differences (percent spliced in, PSI) for all ALS-TE (red), ALS-Ox (blue), and ALS-Glia (gold) transcriptomes to the splicing differences (PSI) for sorted TDP-43-negative cortical neurons^39^. (E) The ALS-Inflam-cord subtype also showed a strong signature for TDP-43 dysfunction, as measured by comparing splicing differences (percent spliced in, PSI) for ALS-Ox-cord (blue), and ALS-Glia-cord (gold) transcriptomes to the splicing differences (PSI) for laser microdissected ALS motor neurons^35^. (F) Altered splicing patterns for the known TDP-43 target gene UNC13B in ALS-TE and ALS-Glia subtype samples from the cortex. (G) Altered splicing patterns for the known TDP-43 target genes STMN2 and UNC13B in the ALS-Glia-cord samples.

We next explored correlations between TDP-43 dysfunction and ALS subtypes in the cortex and spinal cord. Previous analysis of the ALS cortex data had shown strong enrichment for dysregulation of TDP-43 target genes among the set of markers for the ALS-TE subtype in the cortex and an enrichment for dense TDP-43 pathology in IHC stains of tissues from that group^1^. Yet, no equivalent of the ALS-TE cortical subtype was found in the spinal cord tissues. To better address questions of TDP-43 dysfunction in these samples, we took advantage of a recent study profiling ALS/FTD patients with and without functional TDP-43 protein in the nucleus where it normally resides (Liu et al 2019). Isolating all neuronal nuclei from the cortex, they then counter-sorted for TDP-43 positive and negative cells and performed bulk RNA sequencing of the two sorted populations. Performing differential expression and differential splicing analysis, they identified thousands of differentially expressed and differentially spliced genes that mark cells without functional TDP-43. Later studies incorporating these data have identified hallmark TDP-43 splicing targets such as STMN2^29,30^, UNC13A^31,32^ and others^33^. To compare these differential splicing changes to those seen in the NYGC ALS Consortium data, we used Leafcutter^34^ to identify TDP-43 targets that also show evidence of differential splicing across the subtypes in the spinal cord and cortical tissues (Table S6). Figure 3D shows differential splicing changes plotted as Delta percent spliced in (Delta PSI) for each of the ALS subtypes vs control in the cortex (y-axis) as a function of the Delta PSI values for those same genes in the TDP-43 depleted neurons from Liu et al (2019). Both the ALS-TE and ALS-Glia subtypes showed evidence of significant correlation between the measured splicing changes seen in cortical tissues and those seen in sorted TDP-43 negative neurons (Fig 3D), with the ALS-TE subtype samples showing the strongest correlation in Delta PSI values (Pearson’s r = 0.9, P<1e-4). This includes alterations in known TDP-43 target genes such as UNC13B (Fig 3F). In contrast, the ALS-Ox subtype samples from cortex showed no significant association between splicing changes in these samples and those from sorted TDP-43 negative neurons. These data suggest that both ALS-Glia and ALS-TE patient cortex tissues are showing evidence of TDP-43 dysfunction in the tissue that is strong enough to detect in bulk samples, with ALS-TE samples showing the greatest evidence for TDP-43 dysfunction-mediated splicing alterations. These results mirror the results from our previous ALS subtype study^1^ where ALS-TE subtype tissues had the highest levels of TDP-43 pathology as measured by IHC stains for pTDP-43.

To extend this analysis to the spinal cord samples, we turned to a set of previously published laser micro-dissected spinal motor neurons from ALS and control spinal cord samples^35^. Using this spinal motor neuron reference set, we saw strong correlations between splicing alterations seen in the NYGC Cohort ALS-Glia spinal cord samples (Fig 3E) and the splicing alterations seen in the Krach et al^35^ motor neurons (Pearson’s r = 0.71, P∼0). This includes alterations in known TDP-43 target genes such as UNC13B^32,36^ and STMN2^29,30^ (Fig 3G).

Taken together, this integrated analysis shows significant evidence for TDP-43 mediated splicing defects in both the ALS-TE cortex samples and the ALS-Glia/inflammatory cortex and spinal cord samples. It is unclear why the ALS-Ox subtype tissue samples lacked evidence for strong TDP-43 dysfunction as measured by splicing defects, though it is possible that these defects are blurred in bulk samples by a mixture of cells with varying levels of TDP-43 dysfunction.

### 5. Integrative analysis across spinal cord & cortex reveals correlates of ALS molecular subtypes with disease duration

We next asked whether any ALS subtypes were associated with disease duration. Here, we’ve defined disease duration as the number of months between the date of ALS diagnosis and the date of death or tracheostomy, which varied between 6 and 264 months in our cohort. For all patients with cortical samples (n=277), we plotted the eigengene value for the set of genes that define either the ALS-TE subtype (red, Fig 3B), the ALS-Ox subtype (blue, Fig S4A), or the ALS-Glia/inflammatory subtype (gold, Fig S4B). We then correlated that ALS subtype eigengene value for each patient with disease duration in months. Only the ALS-TE patients showed a strong negative correlation between ALS subtype module eigenvalues and disease duration (r=-0.28, P<0.03). This suggests that elevated expression of the ALS-TE subtype genes in cortex correlates with a shorter disease duration. This is likely equivalent to saying that increased TDP-43 dysfunction in cortex is associated with shortened survival times, since these correlations could largely be recapitulated using a correlation between ALS-TE subtype expression values and differential expression patterns seen in the above-described TDP-43 depleted ALS/FTD neurons (See Supp Fig S3, rho=0.18, P<0.04). No association was seen between expression of the ALS-Ox or ALS-Glia/inflammatory eigengenes and disease duration in cortex despite some correlation between the ALS-Glia subtype and TDP-43 dysfunction (See Supp Fig S4A-B). Given the small number of patients in the ALS-Glia subtype (16%), we may be underpowered to detect a correlation between ALS-Glia subtype signatures and disease duration for this group.

We repeated this exploration of clinical correlates for the spinal cord subtypes. For all patients with spinal cord samples in the NYGC ALS cohort, we looked to see whether expression of the ALS-Glia/inflammatory or ALS-Ox eigengene signature in spinal cord was associated with shortened disease duration. Here, the eigengene associated with the ALS-Glia/inflammatory signature in spinal cord was significantly correlated with shortened disease duration. This was especially true for patients assigned to the ALS-Glia/inflammatory group (Fig 3C, gold, r=-0.25, P<2.3E-04), but was also true for patients assigned to the ALS-Ox cord group (Fig S4C, blue, r=-0.32, P<1.7E-06). This suggests that some patients in the ALS-Ox group in the cord show moderate expression of the inflammatory signature genes (e.g., have some positive eigengene values), and to the extent that they do, this correlates with shortened survival times. Taken together, this argues that inflammatory pathway expression in spinal cord is correlated with shortened disease duration, while it is TDP-43 dysfunction signatures in cortex that is most correlated with shortened disease duration.

### 6. A single-cell adapted deep ALS neural net classifier (scDANCER), shows cell type composition changes among ALS subtypes in cortex

The above analysis was all performed using bulk tissue samples from frontal/motor cortex and spinal cord. These bulk average expression profiles are likely a combination of alterations taking place in specific cell types as well as alterations in the cell type composition of each tissue sample. To explore these possibilities, we obtained single-nucleus RNA-seq (nuc-seq) samples from several patient motor cortex samples with bulk RNA-seq data available. In addition, a large set of ALS motor cortex nuc-seq samples were obtained from an independent cohort^37^. Together, this integrated dataset represents 48 ALS patient nuc-seq transcriptomes (from 26 individuals) as well as 28 control motor cortex samples (from 14 individuals) (Fig 4A). An integrated UMAP projection (Fig 4B) showing all 509,966 cells from the fully integrated dataset shows good representation of excitatory and inhibitory neuronal subtypes expected in the upper cortical layers as well as good representation of astrocytes, microglia, oligodendrocytes, and oligodendrocyte precursor cells (OPCs). Cells in the UMAP projection cluster by cell type first, and not by ALS/control, sex, or data source (Supp Fig S5), indicating good integration of all data. A heatmap of cell-type specific marker genes (Fig 4C) shows faithful expression of cell lineage markers by each corresponding cell type, with a Layer 5 extrathalamic (L5ET) cortical neuron signature represented in this heatmap by the markers FEZF2 and LRATD2^38^, an excitatory neuron subtype that includes cortical motor neurons (Betz cells) as well as other layer 5 cortical excitatory neurons.

**Figure 4:**
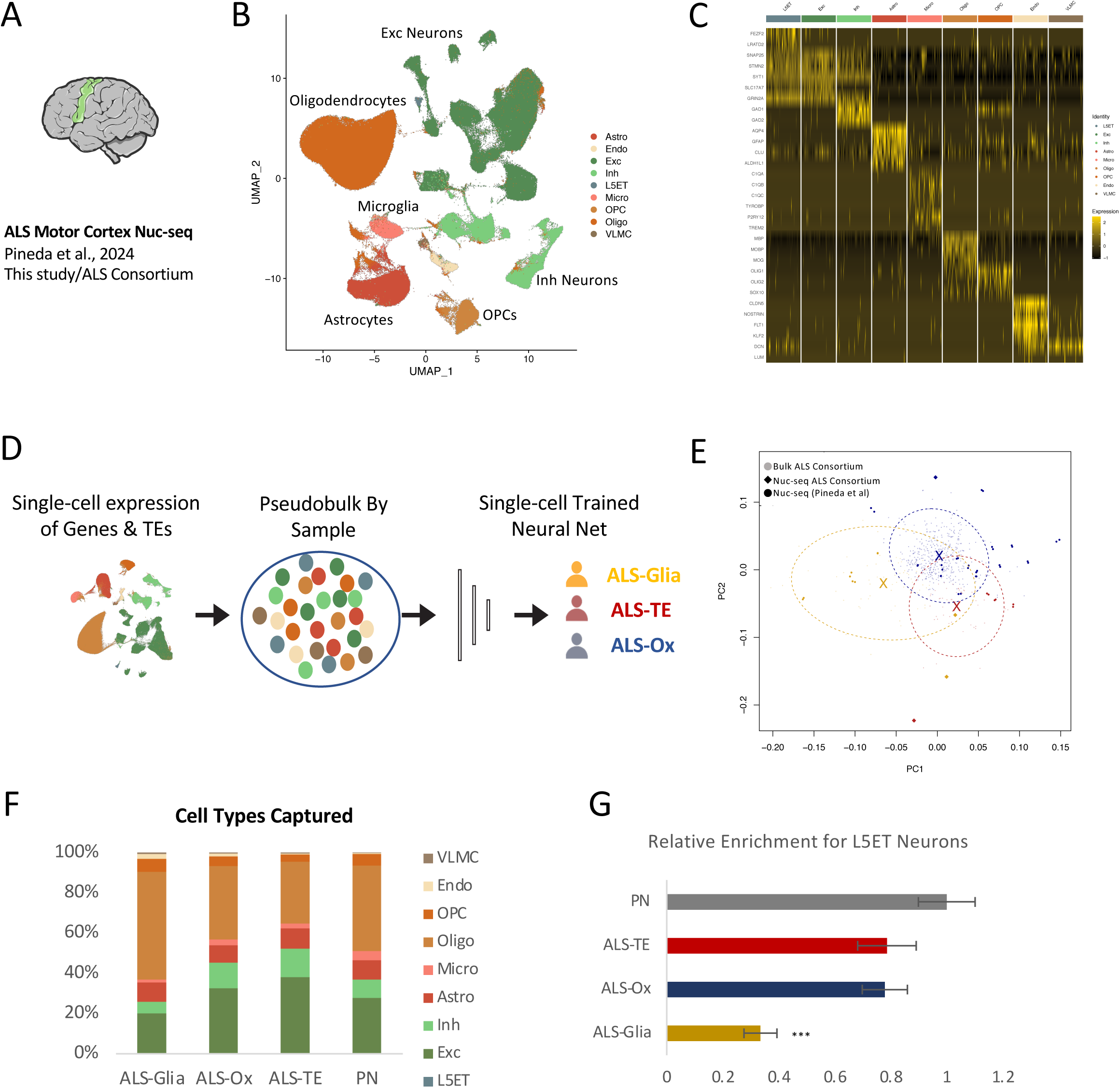
A single-cell adapted ALS neural net classifier (scDANCer) shows cell type composition differences between ALS Subtypes. (A) Single-nucleus RNA-seq (snRNA-seq) transcriptomes were generated for 6 motor cortex samples from the NYGC ALS Consortium, which were integrated with an additional XX ALS and control snRNA-seq motor cortex from Pineda et al. to generate an snRNA-seq ALS motor cortex atlas. (B) A UMAP project of the full integrated snRNA-seq dataset shows good representation of all major cell types in the motor cortex including the L5ET neurons that include motor neurons. (C) A heatmap of cell type specific expression patterns for each of the identified cell types in the motor cortex. (D) A schematic of the adapted snRNA-seq version of our ALS classifer, scDANCer, calls ALS subtype for pseudobulk transcriptomes from snRNA-seq data. (E) Both bulk RNA-seq and pseudobulked snRNA-seq transcriptomes occupy the same region of a PCA plot, which separates samples by ALS subtype and not by data platform (bulk/snRNA-seq) or dataset origin (this study/Pineda et al). Each sample is colored by ALS Subtype (ALS-TE: red, ALS-Ox: blue, ALS-Glia: gold) and ellipses are drawn to encompass 90% of all data from each subtype (F) Across ALS Subtypes, we see that ALS-Ox samples have the same cell type composition as pathologically normal (PN) samples, while ALS-Glia and ALS-TE subtype samples show deviations. (G) For L5ET neurons, which include motor neurons, we see significant differences in the relative fraction of neurons present in ALS-Glia subtype samples as compared to PN controls.

For the samples sequenced for this study, the ALS subtypes of those patient tissues are already known from bulk RNA-seq profiles. For the additional set of published ALS nuc-seq samples^37^, no corresponding bulk RNA-seq data was available to assign an ALS subtype via our trained ALS classifier, DANCer. To overcome this, we retrained our classifier to take in pseudobulk data from the nuc-seq composite transcriptomes and accurately assign ALS subtypes directly from the nuc-seq data (Fig 4D). Specifically, we took the labeled bulk RNA-seq data that had previously been used to train DANCer and down-sampled the data to only select genes detectable in the nuc-seq transcriptomes. This down-sampled version of the classifier reached 91.8% accuracy on test/train data after 200 training epochs, using the same 20-fold cross-validation strategy to estimate classifier accuracy as was used for DANCer (See Fig S6 and Table S7A). We next challenged the single-cell DANCer classifier (scDANCer) with nuc-seq samples obtained from adjacent tissue sections of the same samples used for bulk profiling. scDANCer accurately called the correct ALS subtype for these six samples. Moreover, the pseudobulk transcriptomes could be directly projected onto a PCA plot that includes all bulk NYGC ALS Consortium samples (Fig 4E). On this PCA plot (Fig 4E) the pseudobulk samples are represented by large diamonds for the ALS Consortium nuc-seq samples and large circles for the independent cohort of nuc-seq data from Pineda et al^37^. Small circles represent the bulk NYGC ALS Consortium data. For the NYGC ALS Consortium samples, we confirmed that the scDANCer subtype assignments were consistent with the known subtypes previously obtained from bulk profiles (Supp Table 7B). For the Pineda et al. samples^37^, we confirmed that replicate samples from the same donors gave the same subtype assignments for 95% of the samples (Supp Table 7C).

Once subtype was assigned to all ALS samples in the combined integrated nuc-seq cohorts, we next looked to assess whether cell type composition changes explain some of the transcriptional differences seen across ALS Subtypes. As shown in Fig 4F, we see differences in the number of cells sampled across ALS subtypes. The ALS-Ox samples show a cell type composition that was largely reflective of that seen in pathologically normal motor cortex, with mild but significant enrichments for excitatory (p<0.01) and inhibitory neurons (p<0.001) and slightly more pronounced depletion of microglia (p<0.0002) captured. Similarly, the ALS-TE samples show a slight enrichment for excitatory (p<0.01) and inhibitory neurons (p<0.01) and relative depletion of oligodendrocyte precursor cells (p<0.006) and microglia (p<0.002) captured. Conversely, the ALS-Glia samples show a more pronounced depletion of both excitatory (p<0.006) and inhibitory neurons (p<0.01) among the cells captured, and a relative increase in microglia (p<0.01). Looking more specifically at just the L5ET neuron populations, all ALS patient samples showed fewer L5ET neurons captured relative to those present in pathologically normal samples, likely reflecting an increase in L5ET neuron loss across all ALS samples, which includes motor neurons (Figure 4G). However, only the ALS-Glia/inflammatory subtype samples show both a substantial and significant loss of L5ET neurons, with 70% fewer motor neurons recovered, as compared to controls (P<6E-06).

### 7. ALS L5ET neuron profiles show both pan-ALS and ALS subtype-specific alterations

We next set out to determine whether all ALS L5ET neurons would show similar alterations in gene expression, or whether, conversely, differences would emerge between L5ET neurons from tissues with different ALS subtypes. We first identified all differentially expressed genes between ALS and control L5ET neurons, finding hundreds to thousands of differentially expressed (DE) genes in each ALS subtype (Fig 5A, Table S8). While splicing analysis is difficult in short-read nuc-seq data, we can infer relative levels of TDP-43 dysfunction in these L5ET neurons using gene set enrichment analysis (GSEA) based upon the set of genes known to be down-regulated in sorted TDP-43 negative cortical neurons^39^ as described above. Figure 5B shows the GSEA calculated normalized enrichment score (NES) and adjusted P-values for TDP-43 down-regulated genes in L5ET neurons from each of the ALS subtypes. The ALS-TE subtype L5ET neurons showed the greatest evidence for TDP-43 associated gene expression changes out of all ALS L5ET neurons (NES=-2.1, FDR<0.003) while L5ET neurons from the other subtypes failed to show significant alteration (Log_10_(FDR)>1.3) in the gene set identified for TDP-43 negative neurons^39^.

**Figure 5:**
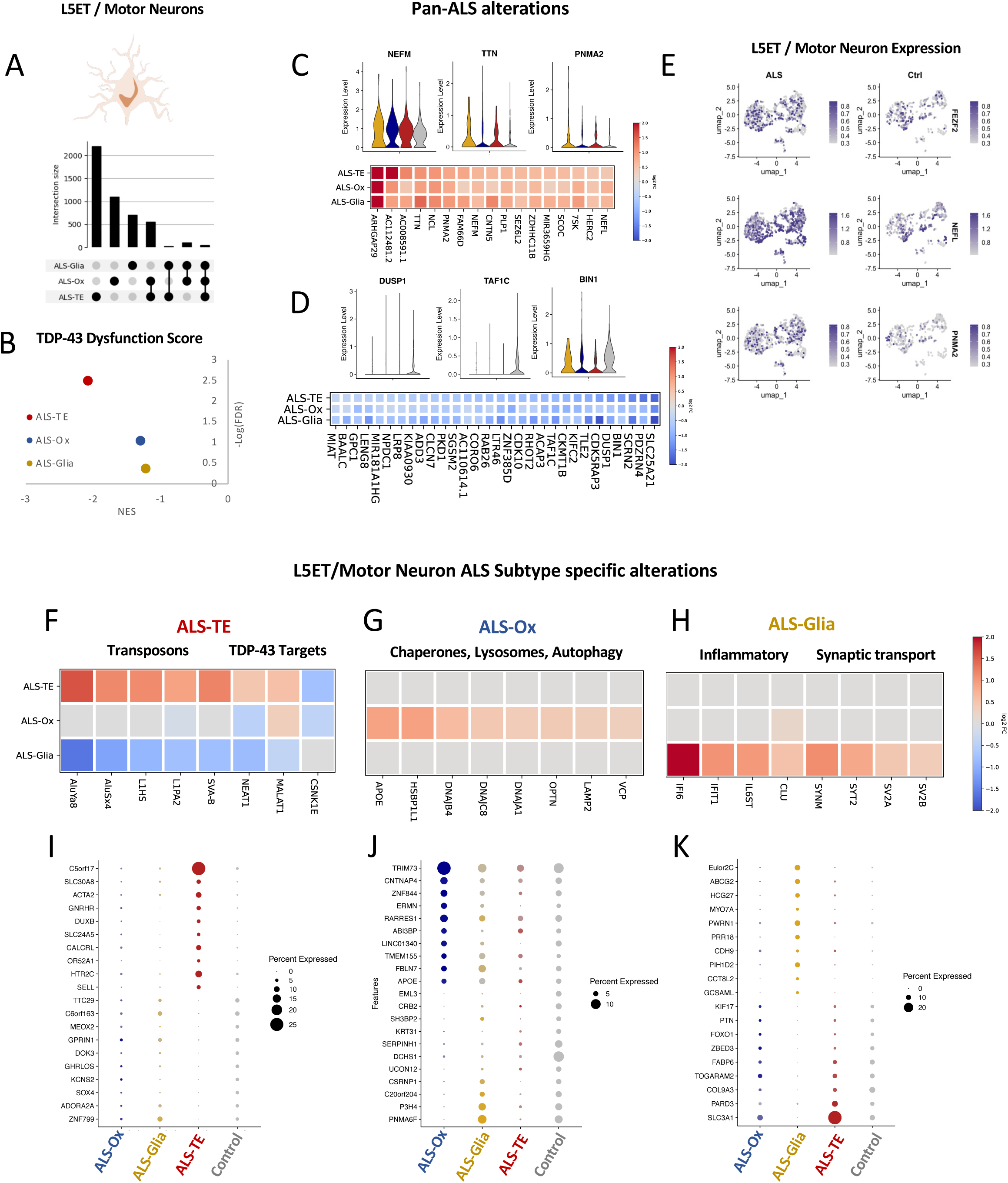
ALS L5ET single neuron expression profiles show both ALS-wide and ALS subtype-specific alterations. (A) We compared the significantly differentially expressed (DE) genes in L5ET neurons between each ALS subtype and PN control samples. While each ALS subtype showed thousands of DE genes, very few of these genes overlapped across subtypes, as shown by the upset plot. (B) To determine whether these L5ET neurons showed signs of TDP-43 dysfunction, we used GSEA to determine whether each subtype showed significant alterations in known TDP-43 target genes (identified from FACS sorted TDP-43 negative neurons^39^). Those GSEA enrichment score (NES) are plotted against the enrichment P-values (plotted as Log(FDR)). Only L5ET neurons from ALS-TE subtype samples show significant alterations in TDP-43 target expression. (C) Shared up-regulated genes across all ALS subtype L5ET neurons are shown as a heatmap of fold-change values, with violin plots for selected genes (NEFM, TTN, PNMA2). (D) Shared down-regulated genes across all ALS subtype L5ET neurons are shown as a heatmap of fold-change values, with violin plots for selected genes (DUSP1, TAF1C, BIN1). (E) A heatmap of gene expression is overlayed for all L5ET cells in the UMAP projection for the motor neuron lineage gene FEZF2 as well as selected pan-ALS down-regulated genes (NEFL, PNMA2). (F-H) A heatmap of fold change values is shown for selected ALS subtype specific alterations in L5ET neurons that were also shown to be subtype-specific in bulk ALS subtype samples. (I-K) The top 10 most ALS subtype specific up- and down-regulated genes is shown as a dot plot, where dot size corresponds to the percent expression of that gene in L5ET neurons of each subtype for ALS-TE (I), ALS-Ox (J), and ALS-Glia(K) samples.

While the results in Figures 5A-B highlight differences between the ALS subtypes in terms of L5ET expression, there were shared alterations, or pan-ALS L5ET neuronal expression changes. Looking across all L5ET neurons in the ALS samples as compared to controls, we found 17 shared genes significantly up-regulated and 30 genes significantly down-regulated across all ALS subtype L5ET neurons (Fig 5B, Table S8G). These shared alterations largely reflect neuronal stress genes that have previously been associated with ALS (NEFL, NEFM), pro-apoptotic genes (PNMA2, BIN1), and genes generally involved in proteostasis (DUSP1, SCRN2). Differential expression heatmaps and violin plots of these pan-ALS genes is shown for all ALS sample and control L5ET neurons in Fig 5C-D. UMAP displays of gene expression in ALS and control L5ET neurons for a subset of these pan-ALS altered genes are shown in Fig 5E below the L5ET marker gene FEZF2.

In addition to these shared ALS-wide expression alterations, L5ET neurons also displayed subtype-specific expression alterations reflective of gene pathways previously identified in the bulk tissue profiles (See Table S8). L5ET neurons of the ALS-TE subtype show broad up-regulation of transposable element (TE) transcripts from young and active TEs (Fig 5F), such as the human specific LINE-1 element (L1HS), and young Alu TEs (AluYa8, AluSx4). These ALS-TE L5ET neurons also show dysregulation of known targets of TDP-43, such as the long non-coding RNAs NEAT1^40,41^ and MALAT1^40,41^ as well as the TDP-43 kinase CSNK1E^35^ (Fig 5F). In addition to these selected genes that overlap with those previously identified in bulk profiles, we show the top 10 most subtype-selective differentially expressed up- and down-regulated genes in ALS-TE L5ET neurons (Fig 5I).

L5ET neurons from ALS-Ox subtype motor cortex show broad up-regulation of genes associated with autophagy, lysosomes, and general chaperone function (Fig 5G). This includes the ALS-associated genes VCP and OPTN, the Alzheimer’s disease associated gene APOE, and several chaperones (HSBP1L1, DNAJB4, DNAJC8). In addition to these selected genes that overlap with those previously identified in bulk profiles, we show the top 10 most subtype-selective differentially expressed up- and down-regulated genes in ALS-Ox L5ET neurons (Fig 5J).

Motor/L5ET neurons from ALS-Glia/inflammatory subtype motor cortex show broad up-regulation of genes associated with inflammatory pathways and synaptic transport (Fig 5H). This includes the interferon induced transcripts (IFI6 and IFIT1), an Alzheimer’s Disease associated apolipoprotein (CLU), and multiple genes associated with synaptic vesicle function and transport (SYT2, SV2A, SV2B). In addition to these selected genes that overlap with those previously identified in bulk ALS-Glia profiles, we show the top 10 most subtype-selective differentially expressed up- and down-regulated genes in ALS-Glia L5ET neurons (Fig 5K).

### 8. ALS subtype-specific differences in pro-inflammatory pathway activation in astrocytes

Given that astrocytes play central roles in mediating neuroinflammatory processes in the central nervous system^42^, we next sought to understand whether these cells would also show ALS subtype-specific differences in their expression patterns. To explore this possibility, we extracted astrocytes from all samples and reclustered just the captured astrocytes, with a UMAP projection of their expression in figure 6A. We used Leiden clustering^43^ to determine whether there might be regions of the UMAP projection space that reflect different functional states. Leiden returned 14 clusters, 9 of which were present in a single cluster without evidence of doublets or cell type assignment errors (Supp Table S9A). Looking at the genes that marked each of these astrocyte clusters (Sub Table S9B) and comparing to previously published studies of astrocyte markers^25,26,44^, we noticed that the 9 high confidence astrocyte clusters could be broadly grouped into 4 main expression programs, which are labeled on the UMAP representation (Fig 6a): those expressing homeostatic markers of astrocyte function and marked by genes such as WIF1 (Fig 6b, top left) those expressing interferon associated gene pathways (cite) and marked by high CD44 expression (Fig 6b, top right), genes that have been previously identified as marking Disease Associated Astrocytes (DAA)^25^ and show high expression of TMSB4X (Fig 6b, bottom left) and those that express high levels of TEs and the receptor P2RY14 (Fig 6b, bottom right).

**Figure 6:**
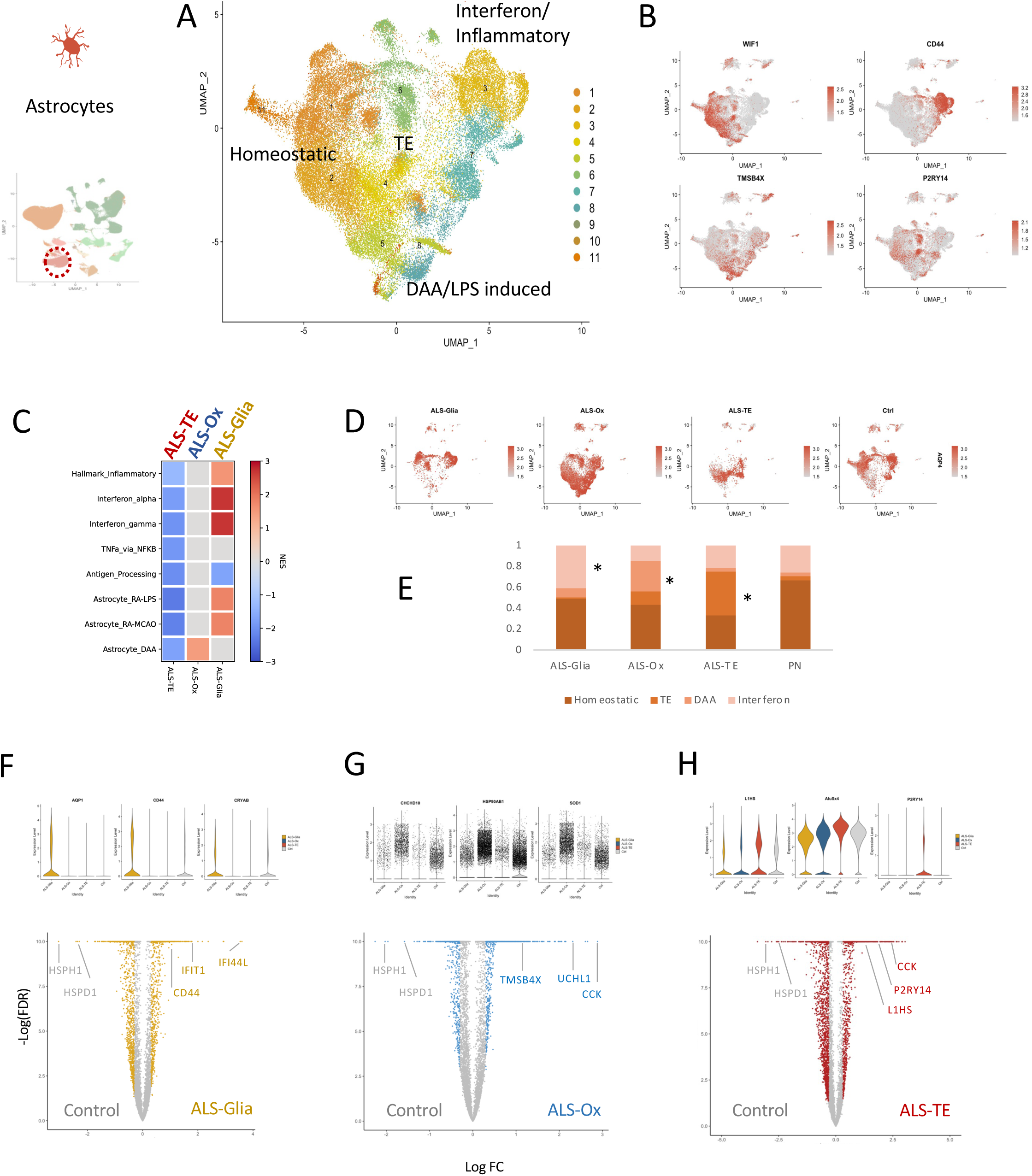
ALS subtypes show differences in pro-inflammatory pathway activation in astrocytes. (A) Astrocytes from all ALS and PN samples were selected and re-clustered using Leiden clustering. The gene markers that identify each Leiden cluster overlapped for many identified clusters, allowing for a broad grouping of astrocytes into 4 main categories marked by genes associated with: interferon-driven inflammatory signaling, LPS induced disease-associated-astrocytes (DAA), basic homeostatic astrocyte function, and transposable elements (TEs). (B) For each of these broad categories, a heatmap of marker gene expression is overlayed on the UMAP projection plot, including the homeostatic marker WIF1, the pro-inflammatory marker CD44, the DAA marker gene TMSB4X, and the purine receptor P2RY14. (C) GSEA enrichment scores (NES) are plotted for several published astrocyte pro-inflammatory markers across ALS subtype astrocytes. The ALS-Glia subtype astrocytes showed enriched expression of interferon pathway genes, markers of astrocytes responding to LPS stimulation (RA-LPS), and markers of oxidative stress induced reactive astrocytes (RA-MCAO). The ALS-Ox subtype astrocytes showed enrichment for expression of “disease associated astrocyte” (DAA) markers previously identified from Alzheimer’s disease models. ALS-TE astrocytes showed no significant enrichment for these inflammatory markers. (D) Astrocytes from each ALS subtype occupy different regions of the UMAP projection space, as shown by plotting the expression of a pan-astrocyte marker, AQP4. (E) Differential occupancy of the UMAP projection space also corresponds to differential representation in the broad astrocyte groupings described above, with ALS-Glia astrocytes significantly enriched in the interferon-marked group, ALS-Ox astrocytes significantly enriched in the DAA-marked group, ALS-TE astrocytes significantly enriched in the TE-expressing group, and PN control astrocytes predominantly composed of cells expressing homeostatic markers. (F) Comparing gene expression for all ALS-Glia astrocytes to controls identified several ALS-Glia specific DE genes (AQP1, CD44, CRYAB, IFI44L) as well as down-regulated genes shared by all ALS astrocytes (HSPH1, HSPD1). (G) Comparing gene expression for all ALS-Ox astrocytes to controls identified several ALS-Ox specific DE genes (CHCHD10, HSP90AB1, SOD1, UCHL1) as well as down-regulated genes shared by all ALS astrocytes (HSPH1, HSPD1). (H) Comparing gene expression for all ALS-TE astrocytes to controls identified several ALS-TE specific DE genes (L1HS, AluSx4, P2RY14) as well as down-regulated genes shared by all ALS astrocytes (HSPH1, HSPD1) and a gene shared with ALS-Ox astrocytes, CCK.

We next explored whether these broad astrocyte clusters could be associated with ALS subtypes, and vice versa. Starting from the ALS subtype associated changes, fold changes for all detected genes in astrocytes of a given ALS subtype were used as input for Gene Set Enrichment Analysis (GSEA)^24^ using a modified MSigDB database^24^ that included KEGG and Hallmark Pathways (see Methods). In addition, we combed the literature for gene sets representing the transcriptional response of astrocytes to particular pro-inflammatory signals. This includes reactive astrocytes induced by lipopolysaccharide (RA-LPS^26^) reactive astrocytes induced by oxidative stress (RA-MCAO^26^), and disease-associated astrocytes (DAA^25^). We expected that we might see elevated pro-inflammatory pathway expression in astrocytes from ALS-Glia/inflammatory subtype samples. Consistent with this, ALS-Glia/inflammatory astrocytes showed significant and substantial upregulation of genes in the several pathways associated with an innate immune response: Interferon-alpha, Interferon-gamma, TNF-alpha mediated NFKB signaling, and the Hallmark Inflammatory pathways (Fig 6c, Table S9A, S9C-D). There was no significant alteration of adaptive immune system pathways, such as the Antigen Processing and Presentation gene set (Fig 6c). Of the experimentally-defined pro-inflammatory astrocyte gene sets, genes induced by both LPS stimulation (RA-LPS) and oxidative stress (RA-MCAO) were elevated in ALS-Glia/inflammatory astrocytes (Table S9D). We were surprised to find that only the ALS-Ox subtype astrocytes showed elevation of genes associated with the “disease associated astrocyte” (DAA) signature, which was not enriched in ALS-Glia astrocytes. The ALS-TE subtype astrocytes did not show enrichment for previously described pro-inflammatory astrocyte pathways.

Using the pseudobulked GSEA analysis alone, we could not determine whether these subtype-specific expression pathway enrichments could be present in all astrocytes from samples of a given ALS subtype. Conversely, these overall enrichments might be a reflection of differential distribution of cells from the above defined astrocyte clusters (homeostatic, TE, DAA, and interferon) in each ALS subtype population. To explore this, we split the UMAP representations to reflect astrocytes of each individual ALS subtype, using a pan-astrocyte marker (AQP4) in Fig 6D. We noticed that ALS-Glia subtype samples contained more astrocytes of the interferon-driven inflammatory subtype (Fig 6D) and that this was significantly different than the distribution of astrocytes in control samples (Fig 6E). Similarly, the ALS-Ox subtype samples were enriched for astrocytes expressing genes previously marking “disease associated astrocyte” clusters (DAA) and those induced by LPS stimulation (Fig 6d), which was specifically enriched in the ALS-Ox samples relative to controls (Fig 6e). Finally, the ALS-TE subtype samples did not show enrichment for astrocytes from the clusters that express inflammatory signals of interferon pathways or DAA marker genes, but did show elevated TE expression (Fig 6d), which was largely absent from controls (Fig 6e).

To further examine these ALS subtype specific differences in astrocyte expression, we identified all genes that specifically mark astrocytes of each ALS subtype as different from each other and from control samples. ALS-Glia astrocytes show significant elevation of genes such as AQP1, CD44, and CRYAB, as shown in violin plots (Fig 6F) and also show interferon induced transcripts (IFIT1, IFI44L) as the top most up-regulated genes in volcano plots (Fig 6F). ALS-Glia astrocytes show down-regulation of several heat shock proteins, such as HSPH1 and HSPD1, which are among the most down-regulated genes in ALS-Glia astrocytes (Fig 6F). See Table S9C for a list of all DE genes in ALS-Glia astrocytes. ALS-Ox astrocytes show elevation of known ALS-associated genes^45^, such as CHCHD10, and SOD1 (Fig 6g, violins). The most upregulated genes in ALS-Ox astrocytes include CCK, UCHL1, and the DAA marker gene TMSB4X (Fig 6G, volcano); the most down-regulated genes in ALS-Ox astrocytes include HSPH1 and HSPD1, as was previously seen for ALS-Glia astrocytes. See Table S9E for a list of all DE genes in ALS-Glia astrocytes. ALS-TE astrocytes show significant elevation of young TEs such as L1HS and AluSx4 (Fig 6h, violins) as well as the P2Y receptor P2RY14, which has previously been associated with response to pro-inflammatory signaling in astrocytes^46^. ALS-TE astrocytes share upregulation of the neuropeptide CCK with ALS-Ox astrocytes (Fig 6h, volcano) as well as down-regulation of the heat shock proteins HSPH1 and HSPD1, which seems to be shared by all ALS astrocytes (Fig 6h). See Table S9G for a list of all DE genes in ALS-TE astrocytes and Table S9I for a list of shared alterations common to all ALS subtype astrocytes.

### 9. ALS subtype-specific differences in pro-inflammatory pathway activation in microglia

Microglia are tightly integrated with astrocytes and neurons in neuroinflammatory cascades, adopting a multitude of states in response to neuronal stress and inflammatory stimuli^47^. Using a similar strategy from the above results exploring subtype-specific differences in astrocyte expression, we next determined if each ALS subtype would show subtype-specific differences in microglial expression and microglia cluster representation. As above, we first extracted and reclustered all captured microglia, with a UMAP projection of their expression in Figure 7A. We used Leiden clustering to understand whether there might be regions of the UMAP projection space that might reflect different functional states. Leiden returned 11 clusters, 7 of which were present in a single cluster without evidence of doublets or cell type assignment errors (Supp Table S10A). Looking at the genes that marked each of these microglia clusters (Sub Table S10B) and comparing to previously published studies of microglial state markers^27,28^, we noticed that the microglia clusters could be broadly grouped into 4 main expression programs, which are labeled on the UMAP representation (Fig 7a): genes associated with elevated metabolic activity and marked by genes such as GAPDH (Fig 7b, top left), those expressing homeostatic markers of microglia function and marked by genes such as P2RY12 (Fig 7b, top right) those expressing interferon associated gene pathways and marked by high APOE expression (Fig 7b, bottom right), and finally genes that have been previously associated with phagocytic microglia activity and marked by high expression of SLC11A1 (Fig 7b, bottom left).

**Figure 7:**
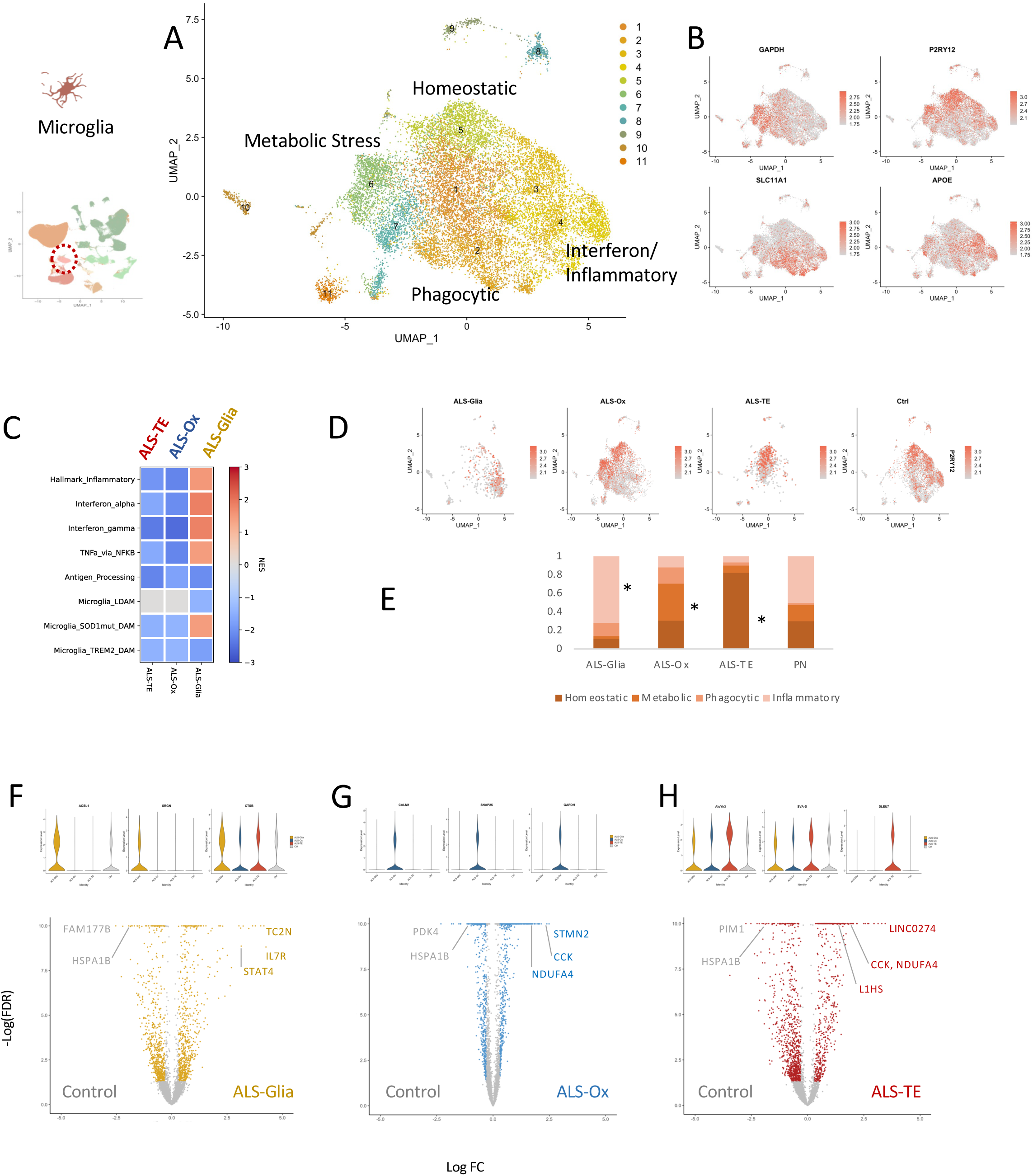
ALS subtypes show differences in pro-inflammatory pathway activation in microglia. (A) Microglia from all ALS and PN samples were selected and re-clustered using Leiden clustering. The gene markers that identify each Leiden cluster overlapped for many identified clusters, allowing for a broad grouping of microglia into 4 main categories marked by genes associated with: interferon-driven inflammatory signaling, phagocytic expression programs, metabolic stress, and basic homeostatic function. (B) For each of these broad categories, a heatmap of marker gene expression is overlayed on the UMAP projection plot, including the metabolic marker GAPDH, the homeostatic marker P2RY12, the apolipoprotein APOE, and the phagocytic marker SLC11A1 (aka NRAMP1). (C) GSEA enrichment scores (NES) are plotted for several published microglia pro-inflammatory markers across ALS subtype microglia. The ALS-Glia subtype microglia showed enriched expression of interferon pathway genes, TNFα/NFKβ pathway genes, and markers identified in “disease associated microglia” isolated from a SOD1 mutant mouse^28^. Neither the ALS-Ox nor the ALS-TE microglia showed significant enrichment for these inflammatory markers. (D) Astrocytes from each ALS subtype occupy different regions of the UMAP projection space, as shown by plotting the expression of a pan-microglia marker, P2RY12. (E) Differential occupancy of the UMAP projection space also corresponds to differential representation in the broad microglia groupings described above, with ALS-Glia microglia significantly enriched in the inflammatory group, ALS-Ox microglia significantly enriched in the metabolic stress group, ALS-TE astrocytes predominantly expressing homeostatic markers, and PN control astrocytes predominantly composed of cells expressing either homeostatic or inflammatory markers. (F) Comparing gene expression for all ALS-Glia microglia to controls identified several ALS-Glia specific DE genes (ACSL1, SRGN, CTSB, IL7R) as well as down-regulated genes shared by all ALS microglia (HSPA1B). (G) Comparing gene expression for all ALS-Ox microglia to controls identified several ALS-Ox specific DE genes (CALM1, SNAP25, GAPDH, STMN2) as well as down-regulated genes shared by all ALS astrocytes (HSPA1B). (H) Comparing gene expression for all ALS-TE astrocytes to controls identified several ALS-TE specific DE genes (AluYh3, SVA-D, L1HS, DLEU7) as well as down-regulated genes shared by all ALS astrocytes (HSPA1B) and two gene shared with ALS-Ox astrocytes, CCK and NDUFA4.

We next explored whether these broad microglia clusters could be associated with ALS subtypes, and vice versa. Starting from the ALS subtype associated changes, fold changes for all detected genes in microglia of a given ALS subtype (Supp Tables S10C,E,G) were used as input for Gene Set Enrichment Analysis (GSEA) using a modified MSigDB database that included KEGG and Hallmark Pathways (see Methods). In addition, we combed the literature for gene sets representing the transcriptional response of microglia to particular pro-inflammatory signals. This includes pro-inflammatory microglia induced by lipid droplets (Microglia-LDAM)^48^, genes marking pro-inflammatory microglia induced by SOD1 mutations (SOD1-DAM)^28^, and disease-associated microglia induced by constitutive TREM2 activity (TREM2-DAM)^27^. We expected that we might see elevated pro-inflammatory pathway expression in microglia from ALS-Glia/inflammatory subtype samples and indeed, ALS-Glia/inflammatory microglia showed significant and substantial upregulation of genes in the several pathways associated with an innate immune response (Table S10D): Interferon-alpha, Interferon-gamma, TNF-alpha mediated NFKB signaling, and the Hallmark Inflammatory pathways (Fig 7c). There was no significant alteration of adaptive immune system pathways, such as the Antigen Processing and Presentation gene set (Fig 6c). Of the experimentally-defined pro-inflammatory microglia gene sets, we were surprised to find that only the genes associated with SOD1 mutations^28^ were elevated in ALS-Glia/inflammatory microglia. The ALS-Glia/inflammatory subtypes samples did not show elevation of lipid droplet associated microglial markers nor did they show enrichment for TREM2 mutation induced changes. Additionally, neither the ALS-Ox (Supp Table S10F) nor the ALS-TE subtype microglia (Supp Table S10H) showed enrichment for these defined microglial marker sets.

We next split the UMAP representations to reflect microglia of each individual ALS subtype, using a pan-homeostatic microglia marker (P2RY12) in Fig 7D. We noticed striking differences in the representation of microglial clusters among the ALS subtypes. ALS-Glia subtype samples were dominated by microglia from the interferon-driven inflammatory subtype (Fig 7D), a distribution that was significantly different from the distribution of microglia in control samples or other ALS subtypes (Fig 7E). Microglia from ALS-Ox subtype samples showed significantly elevated representation from the microglia cluster expressing genes marking elevated metabolic activity (Fig 7d-e). Surprisingly, the ALS-TE subtype samples showed strong enrichment for microglia from the homeostatic cluster (Fig 7d), which was enriched even above the representation of homeostatic cluster microglia in the pathologically normal controls (Fig 7e).

In addition to these cluster representation analyses, we identified all genes that specifically mark microglia from each ALS subtype as different from the other subtypes and from control samples. ALS-Glia microglia show significant elevation of pro-inflammatory marker genes such as ACSL1, SRGN, and CTSB, as shown in violin plots (Fig 7F) and also show elevation of innate immune pathway markers (TC2N, IL7R, STAT4) as the top most up-regulated genes in volcano plots (Fig 6F). ALS-Ox astrocytes show elevation of known ALS-associated genes^5^, such as CHCHD10, HSP90AB1, and SOD1 (Fig 6g, violins). The most upregulated genes in ALS-Ox microglia include CALM1, SNAP25, and GAPDH (Fig 6G, violins) as well as the TDP-43 target gene STMN2 (Fig 6G, volcano). ALS-TE microglia show significant elevation of young TEs such as SVA-D and AluYh3 (Fig 7h, violins) as well as the youngest human transposon L1HS (Fig 7h, volcano).

Similar to our results for astrocytes, there were heat shock proteins that came up as among the most down-regulated genes across all ALS subtypes, such as HSPA1B, as seen in the volcano plots of Figure 7F-H. Moreover, both ALS-TE and ALS-Ox microglia showed upregulation of the neuropeptide CCK (Fig 7G-H) which was previously seen to be elevated in ALS-Ox astrocytes (Fig 6G) as well as a mitochondrial complex associated factor, NDUFA4 (Fig 7G-H). A full table of shared and subtype-specific differentially expressed genes is shown in Table S10I.

Neither microglia nor astrocytes of the ALS-TE subtype showed significant elevation of classical pro-inflammatory pathway genes or specific disease associated astrocyte or microglia signatures (DAA/DAM). This suggests that classical interferon signaling pathways may not be active in these cells. However, looking more closely at the lists of differentially expressed gene sets in the ALS-TE microglia and astrocytes, we noticed ALS-TE specific elevation of several genes typically elevated in response to activation of the NLRP inflammasome. This included STAT1 and IL18, key downstream effectors of inflammasome activation^49^. These genes, as well as several interferon induced genes (IFIT2, IFIT3) were specifically elevated in ALS-TE microglia (Table S10G). Elevation of these specific inflammatory genes was coincident with elevation of several young transposable elements (L1HS, AluYc3, SVA-A) in microglia, which were among the most elevated genes across ALS-TE subtype microglia (Table S10G).

## Discussion

In this study, we present an analysis of the largest ALS patient molecular profiling study available, with whole genomes and whole transcriptomes available for hundreds of ALS patients from multiple affected tissues. This large patient cohort integrated across cortex and spinal cord allowed us to explore the molecular landscape of alterations in ALS patient tissues. The size of this study allowed us to conclude that ALS molecular subtypes are highly robust across multiple clinical centers in large patient cohorts. We used *de novo* pattern finding methods to determine that no new molecular subtypes were present in this larger patient cohort that quadrupled the size of the patient samples available in our initial study^1^. ALS molecular subtypes have now been replicated in two independent studies^13,14^, although we note that only one of these studies^13^ represented an independent patient cohort with novel findings^13^.

The availability of previously labeled samples from our original ALS molecular subtype discovery allowed us to develop a highly accurate deep ALS neural net classifier (DANCer) to assign molecular subtypes across tissues in both bulk and single-cell data. DANCer achieved 96% accuracy on previously labeled samples, with the remaining poorly-classified samples likely due to mixing of biological patterns within a single tissue rather than inaccuracy of the classifier itself, as determined by reference to samples with ALS-TE/ALS-Glia mixed calls that, upon IHC stains for pTDP-43 pathology and IBA1-positive microglia, showed evidence consistent with both microglia activation and dense pTDP-43 pathology.

Upon integrating the cortex and spinal cord data available in the updated NYGC ALS Consortium cohort, we noted that ALS subtype patterns in the spinal cord largely mirror those seen in the motor cortex. Patients with an ALS-Glia/inflammatory subtype in the motor cortex are more likely to show inflammatory patterns in their spinal cord samples, as well, suggesting systemic neuroinflammation. The TDP-43 driven ALS-TE subtype does not appear in the bulk spinal cord tissue samples, which may reflect differences in cellular composition of the two tissues. In particular, we note that splicing changes in the ALS-TE samples showed the greatest correlation with previously noted TDP-43 splicing changes in cortex samples, as determined from sorted TDP-43 negative cortical neurons^39^. While the ALS-TE subtype is not present in spinal cord samples, the ALS-Glia subtype spinal cord samples showed strong correlation with TDP-43 associated splicing alterations, as measured by comparing to laser micro-dissected lower motor neurons from ALS spinal cord samples^35^.

This large integrated cortex & spinal cord dataset enabled us to determine clinical correlates of ALS molecular subtypes. Strong correlations with disease duration were seen for ALS-TE subtype expression in the cortex and ALS-Glia subtype expression in spinal cord. Taken together, these results suggest that disease duration in ALS reflects, in part, the degree of TDP-43 dysfunction in the cortex and inflammatory processes in the spinal cord.

After training DANCer on bulk transcriptome data, we next built a single-cell version of the classifier that down-sampled the original bulk samples to the depth and detectability of single-cell data. The classifier remained highly accurate (92%) on pseudo-bulked single-cell data, allowing us to accurately identify ALS molecular subtypes from single-cell data alone, without the need for matched bulk transcriptomes – as measured by samples where we obtained both bulk and single-cell transcriptome data. This single-cell classifier, scDANCer, enabled the incorporation and integration of 48 total ALS single-nucleus transcriptomes from cortex. Single-nucleus profiles show that ALS subtypes are a combination of cellular composition as well as subtype-specific differences in cellular states. The ALS-Glia samples showed the most substantial and significant loss of L5ET neurons, likely reflecting motor neuron loss in ALS. ALS-TE samples did not show significant loss of the L5ET neuron population, but did show the strongest signatures of TDP-43 dysfunction, perhaps reflecting cells that have not yet succumbed to the impact of impaired TDP-43 function.

Of the experimentally-defined pro-inflammatory astrocyte pathways, the ALS-Glia and ALS-Ox samples showed differential enrichment for different inflammatory response markers. ALS-Glia astrocytes showed strong expression of interferon-driven pathways and those that mark response to LPS stimulation^26^ and oxidative stress challenge^26^. In contrast, ALS-Ox subtype astrocytes showed expression of pathways previously annotated as “disease associated astrocytes” in models of Alzheimer’s Disease^25^. Finally, there was a large set of astrocytes that showed elevated TE expression in the absence of clear inflammatory pathway expression, with unknown triggers for that TE expression and unknown consequences for astrocyte function.

Of the experimentally-defined pro-inflammatory microglial pathways, only the SOD1-DAM microglia gene set^28^ showed significant elevation in ALS-Glia/inflammatory subtype microglia (Fig 6a), despite the fact that none of these samples carried known pathogenic SOD1 mutations. This suggests that the SOD1-DAM identified genes from SOD1 mutant mice^28^ may carry information about neuroinflammatory processes in ALS models beyond a simple response to SOD1 mutations. Neither the ALS-TE nor the ALS-Ox subtype microglia showed significant enrichment for classical pro-inflammatory pathways or experimentally defined microglial inflammatory response genes. Taken together these results suggest that different pro-inflammatory pathway genes might be elevated in each of the ALS subtypes, possibly due to differential pro-inflammatory triggers or differential response to inflammatory triggers.

Altogether, the results of this study allowed us to conclude that ALS molecular subtypes are robust across large patient cohorts and replicated across independent patient cohorts^13^. Two of the ALS molecular subtypes are detectable in both motor cortex and spinal cord samples (ALS-Ox and ALS-Glia) with very similar gene expression changes and TDP-43 associated splicing alterations in both tissues. We note that the ALS-Glia subtype was also detected in a previous study of ALS spinal cord samples^16^ and that both studies identified clinical correlates of ALS molecular subtypes with disease duration. Single-nucleus motor cortex transcriptomes enabled us to identify the extent to which cellular composition contributes to bulk molecular subtypes, with the greatest neuronal loss seen in ALS-Glia samples. Splicing analysis enabled us to determine the extent to which ALS molecular subtypes reflect TDP-43 pathology, with ALS-TE and ALS-Glia subtypes both showing evidence for TDP-43 target dysregulation. Thus, ALS molecular subtypes are a combination of pathological changes in TDP-43 function, alterations in cellular composition that include neuronal loss, and ALS subtype-specific and cell-type-specific molecular pathway alterations.

## Methods

### ALS Post-mortem samples

The NYGC ALS Consortium samples in this study were acquired through various IRB protocols from member sites and transferred to NYGC in accordance with all applicable foreign, domestic, federal, state, and local laws and regulations for processing, sequencing, and analyses. All available de-identified clinical and pathological records were collected and used together with C9orf72 genotypes to summarize patient demographics and disease features (see Table S1).

### Generation of bulk RNA-seq data

RNA was extracted from flash-frozen patient samples homogenized in Trizol (15596026, ThermoFisher Scientific, Waltham, MA, USA) -Chloroform and purified using the QIAGEN RNeasy Mini kit (74104, QIAGEN, Germantown, MD, USA). RNA was assessed using the Bioanalyzer (G2939BA, Agilent, Santa Clara, CA, USA). RNA-seq libraries were prepared from 500 ng of total RNA using the KAPA Stranded RNA-seq kit with RiboErase (07962304001, Kapa Biosystems, Wilmington, MA, USA) for rRNA depletion and Illumina compatible indexes (NEXTflex RNA-seq Barcodes, NOVA-512915, PerkinElmer, Waltham, MA, USA). Pooled libraries (average insert size: 375bp) were sequenced on an Illumina HiSeq 2500 or NextSeq V1 using a paired end 125 nucleotide setting, to yield 40-50 million reads per library.

### Analysis of bulk RNA-seq data

Reads from samples with RIN > = 5.5 were aligned to the hg38 human genome using STAR v2.7.6^50^, allowing for a 4% mismatch rate and up to 100 alignments per read to ensure capture of young transposon sequences. Abundance of gene and transposon sequences was calculated with TEtranscripts v2.2.3^51^. For differential expression analysis, we employed DESeq2^52^, using the DESeq normalization strategy and negative binomial modeling. B-H corrected FDR P value threshold of p < 0.05 was used to determine significance. For heatmap visualization, the reads were normalized using a variance stabilizing transformation in DESeq2. Alternative splicing was performed using Leafcutter^34^ following the procedure as outlined in their documentation (http://davidaknowles.github.io/leafcutter/articles/Usage.html)

### Extraction of nuclei from fresh frozen tissue

Single nuclei were extracted from archived human postmortem motor cortex tissue using a method adapted with modifications from Maitra et al^53^. A Dounce homogenizer was used to homogenize fresh frozen cortex tissue on ice. Five mL of chilled Nucleus wash buffer^53^ was added to the homogenate, to quench the lysis. A 30-μm MACs SmartStrainer (Miltenyi Biotech cat. # 130-098-458) was used to remove cell debris. The lysed, filtered homogenate was then promptly centrifuged in a desktop centrifuge equipped with a swinging bucket rotor at 500 G for 5 min at 4°C. After centrifugation, the sample was removed and supernatant decanted without disrupting the nuclei pellet, and placed on ice. The pelleted nuclei were gently resuspended on ice. A fresh 15-mL centrifuge tube was equipped with a 30 μM MACs SmartStrainer and the filtration and resuspension and wash processes were repeated as above. One mL of 50% (weight/volume) working solution of iodixanol (Optiprep) was added to the resuspended nuclei and mixed gently, to obtain 2 mL of nuclei in 25% (wt/vol) iodixanol solution. For gradient centrifugation, 0.5 mL of the nucleus suspension was gently pipetted on top of an iodixanol cushion (0.5 mL of 29% (wt/vol) iodixanol) by letting it slowly run down the wall of the tube, to prevent mixing. Without disturbing the layers, the LoBind tubes containing the nuclei on top of the iodixanol cushion were centrifuged at 10,000 G for 30 min at 4°C. A second gradient centrifugation step was carried out at 10,000 G for 10 min at 4°C. Supernatant was then removed, leaving the nuclei in up to 200 μL buffer volume. Pelleted nuclei were then gently resuspended and filtered through a 40 μm FlowMi Tip Strainer (SP Scienceware cat. # H13680-0040) into a 1.5 mL LoBind tube. Extracted nuclei were immediately place on ice.

### Generation of nuc-seq data

All samples were sorted using the Sony SH800 sorter into 0.04% BSA in PBS and counted on a Countess II FL automated cell counter using 1:1 AOPI stain (Nexcelom CS2-0106). Nuclei were pelleted for 5 min at 500 G in a refrigerated swinging bucket centrifuge and resuspended to a target concentration of 1,000 nuclei/μL prior to loading the 10× Chromium chips. Single-cell gene expression libraries were prepared using the Single Cell 3′ Gene Expression kit v3.1 (10× Genomics, #1000268) according to manufacturer’s instructions. Libraries were sequenced on an Illumina Nextseq2000 using 100-cycle kits to a mean depth of ∼35,000 reads per cell.

### Analysis of nuc-seq data (CellRanger-TE)

In addition to the nuc-seq data described above, raw data from single nucleus sequencing of ALS and pathologically normal (PN) samples^37^ (<u>GSE174332</u>) were downloaded from GEO. All libraries were analyzed using 10x Genomics Cell Ranger version 5.0.1^54^, using a GRCh38 reference genome database built with GENCODE v35 gene annotation^55^ and RepeatMasker TE annotation^56^. Libraries with mapping rate <80%, intergenic (non-gene and non-TE) reads >5% and total read count <10 million were excluded from subsequent analyses. Gene expression estimates of each cell were correlated using Spearman correlation with identified cell types from the human M1 10x dataset^57^, with the cell labelled according to the M1 cell type with the highest correlation.

The processed count matrices from Cell Ranger were integrated in Seurat^58^, with PCA and UMAP performed on the combined dataset. Differential analysis between ALS and control cell types was performed using a modified DESeq2 protocol, where size factor was calculated using the scran R package^59^, a gamma-Poisson GLM (glmGamPoi R package^60^) was used to fit the data, and a likelihood ratio test was used for the statistical comparison. Genes and TEs with an FDR < 0.05 were considered differentially expressed.

### Gene Set Enrichment Analysis

Gene set enrichment was performed using GSEA^24^ and a manually curated set of gene pathways. From MSigDB^24^, this included the Hallmark and KEGG database. From the curated literature, we included differentially regulated genes from TDP-43 depleted neurons^39^, KRAB zinc fingers^61^, and several lists of genes dysregulated in astrocytes^25,26^ and microglia^27,28,48^ from diseased models or other pro-inflammatory stimuli. various disease/mutation/treatment-associated astrocyte and microglia. Gene sets with an FDR < 0.05 were considered significantly enriched/depleted.

### Statistical Data Analysis

All enrichments calculations for discrete variables (e.g., subtype and clinical variables, subtype and cell types or cell clusters) used Fisher’s Exact test, with an alpha=0.05 and a Benjamini-Hochberg adjustment for Type I error. All linear correlations used Pearson’s R with an alpha = 0.05 and a Benjamini-Hochberg adjustment for Type I error. Statistics for differential expression of bulk and single-cell data are described in each section and all involve an alpha=0.05 and a Benjamini-Hochberg adjustment for Type I error. Statistics for differential splice junction usage of bulk data are described above, using an alpha=0.05 and a Benjamini-Hochberg adjustment for type I error. Statistics for GSEA of bulk and single-cell data use an alpha=0.05 and a Benjamini-Hochberg adjustment for Type I error.

### Training and validation of Neural Networks

WGCNA^23^ was used to create eigengenes for the ALS bulk cortex data with a power parameter of 9, resulting in 40 modules. One of these modules was determined to be entirely comprised of sex-specific genes and was removed for the purpose of classifier training. Eigenvalues for each ALS sample were calculated for the remaining 39 modules and used as input to DANCer. DANCer is a feedforward multilayer perceptron composed of 39 input nodes, a fully-connected hidden layer of 6 nodes, and a softmax output layer of 3 nodes that report the probability of classification to each subtype. The classifier was built using the Keras library^22^ as a wrapper for TensorFlow version 2^62^. The labeled ALS subtype samples (n=176) from Tam et al.^1^ were split 80/20 for training/testing, respectively. Training proceeded for 200 epochs with a loss function of categorical cross entropy with callbacks for early stopping to prevent overfitting.

scDANCer is a single-cell(nucleus) adapted version of DANCer that follows the same architecture and training regime. Single nucleus transcriptomes were compressed into pseudobulk representations by summing unique molecular identifier (UMI) transcript counts for all detected genes in all cells of a given sample. scDANCer was trained using the same original bulk RNA-seq data used for DANCer, but the list of input genes was restricted to the set of detectable genes in our pseudobulked nuc-seq transcriptomes. The WGCNA network and eigenmodules were recomputed using those pseudobulked nuc-seq transcriptomes, again using a power parameter of 9, which resulted in 16 eigenmodules. The scDANCer architecture involved 16 input nodes, 6 fully-connected inner nodes, and a softmax output layer of 3 nodes. The labeled ALS subtype samples (n=176) from Tam et al.^1^ were split 80/20 for training/testing, respectively. Training proceeded for 200 epochs with a loss function of categorical cross entropy with callbacks for early stopping to prevent overfitting.

## Availability of data and software

NYGC consortium data for bulk cortex and spinal cord is available at (<u>GSE137810</u>). Single nucleus sequencing data for motor cortex samples matched to a subset of previously profiled^1^ bulk cortex samples are available at (GSE271156). DANCer software for ALS subtype classification is available at https://github.com/mhammell-laboratory/DANcer. CellRanger-TE for single-cell analysis with CellRanger using gene & TE annotation is available at https://github.com/mhammell-laboratory/CellRangerTE.

## Acknowledgements

We wish to thank the Target ALS Human Postmortem Tissue Core for providing post-mortem brain and spinal cord samples, the CSHL Single Cell Core Facility and CSHL Sequencing Facility (supported by an NIH Cancer Center support grant 5P30CA045508) for single-cell data generation and sequencing support, and the Harms laboratory for performing repeat-primed PCR to identify C9ORF72 expansions in the Target ALS samples. Schematic images were created with BioRender.com. M.G.H. was supported by grants from the Chan Zuckerberg Initiative (SVCF2018-191863), the Ride For Life Foundation, the Rita Allen Foundation of which M.G.H. is a scholar, and the NIH/NINDS (RF1NS118570 to M.G.H and H.P). HP was supported by grants from the NIH/NINDS (RF1NS118570 to M.G.H and H.P., R01NS118183, and R01NS116350). K.O. acknowledges funding from the National Science Foundation Graduate Research Fellowship and an NIH training grant (2T32GM065094-16). All NYGC ALS Consortium activities are supported by the ALS Association (15-LGCA-234) and the Tow Foundation. The figures were created with the help of BioRender.com.

## Figure Legends

**Figure S1:**
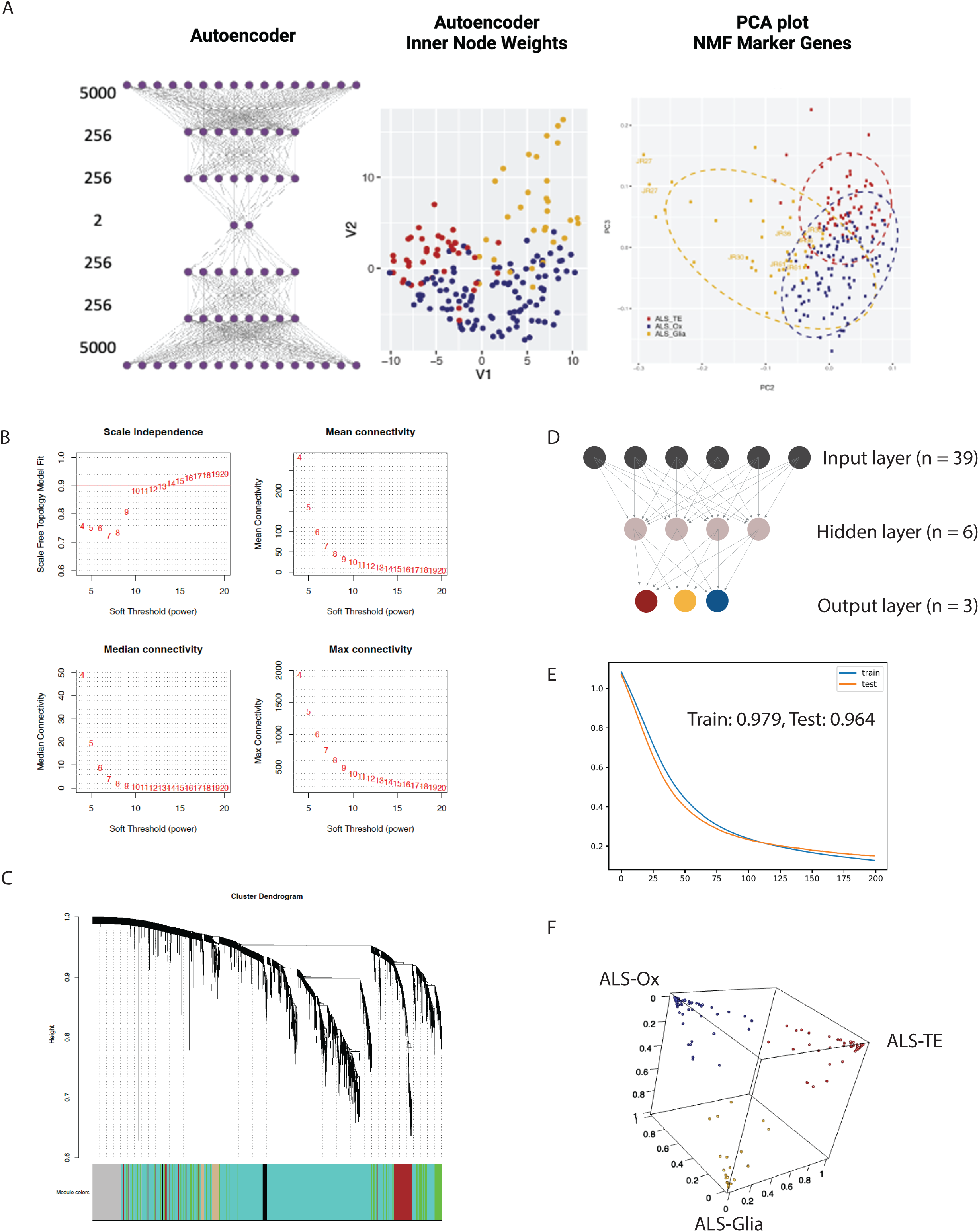
Training of Deep ALS Neural-Net Classifier (DANCer). A) A symmetric autoencoder (left panel: 5000 outer nodes, 2 layers of 256 inner nodes, and a bottleneck layer) was trained to recapitulate the transcriptomes of the ALS motor cortex data from our original discovery dataset^1^. To determine what the autoencoder had learned, we plotted the activation energies for the 2 nodes in the bottleneck layer, also known as the autoencoder latent space (middle panel). The dots for each ALS sample in this autoencoder latent space are colored by that sample’s ALS subtype (ALS-TE: red, ALS-Ox: blue, ALS-Glia: gold). For comparison to the autoencoder latent space, we also show the PCA plot from our original study for those same ALS sample transcriptomes. B) Determining the best soft threshold/power parameter for WGCNA. The power parameter of 9 was chosen due to a scale-free model correlation of ∼0.8 while maintaining connectivity within the network. C) Hierarchical clustering of genes and their assignment to WGCNA modules. D) Layout of the neural net, with 39 nodes in the input layer, 6 nodes in the hidden layer and 3 softmax nodes in the output layer. E) Plot of categorical cross entropy loss in the training and test sets after each epoch of training. The final accuracy is 97.2% for the training set and 96.4% for the test set. F) 3D plot of the classifier confidence from the softmax output, with the dots labeled by their final ALS subtype classification.

**Figure S2:**
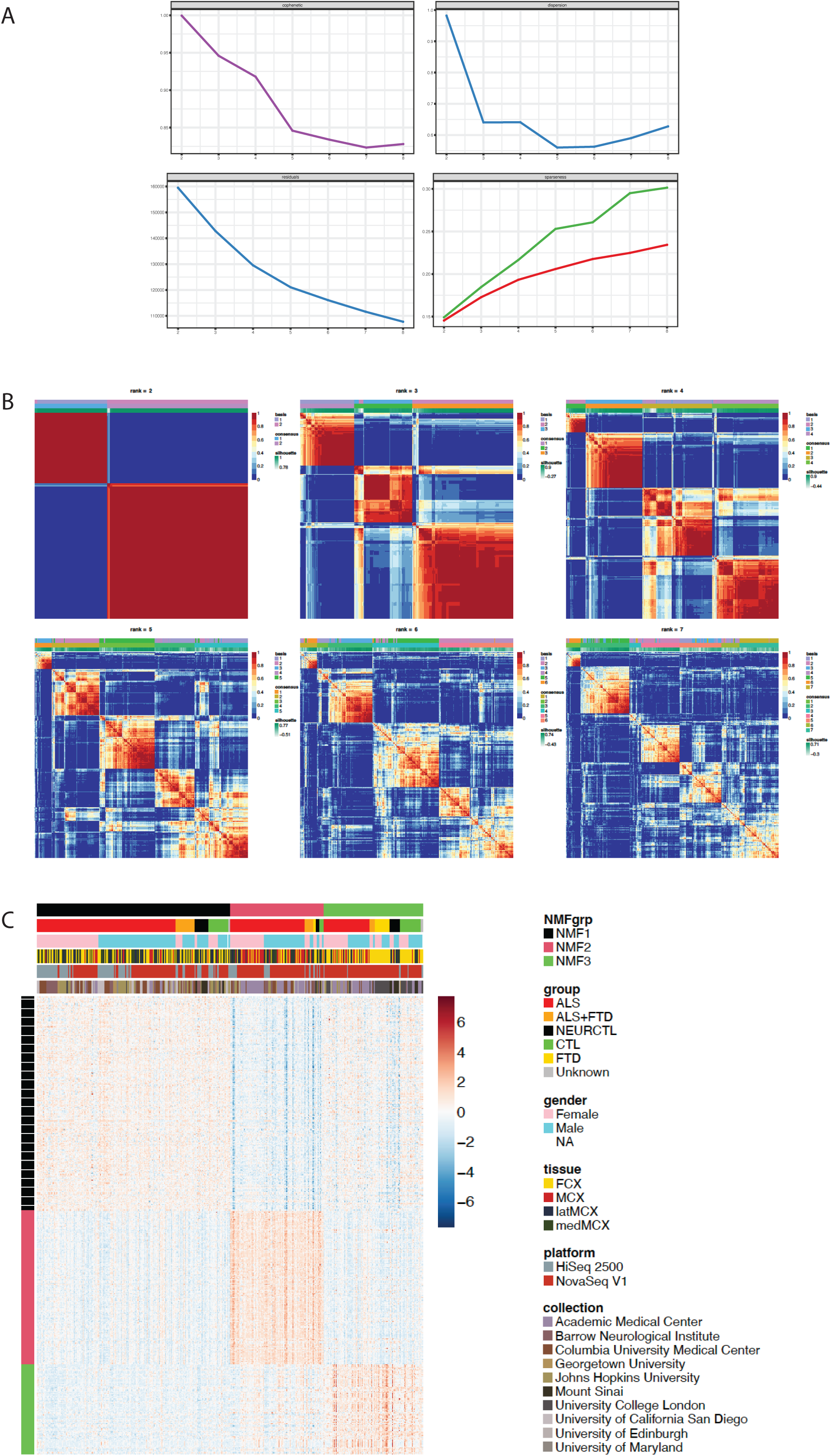
Non-negative matrix factorization of frontal/motor cortex samples. A) Cophenetic (top left), dispersion (top right), residual (bottom left) and sparseness (bottom right) for each rank value tested. B) Consensus map of NMF clusters at rank 2 to 7. Rank of 3 was subsequently chosen due to high cophenetic and low dispersion. C) Expression heatmap of NMF marker genes in frontal/motor cortex samples. NMF assignment does not show obvious correlation with gender, tissue type, sequencing platform or collection site.

**Figure S3:**
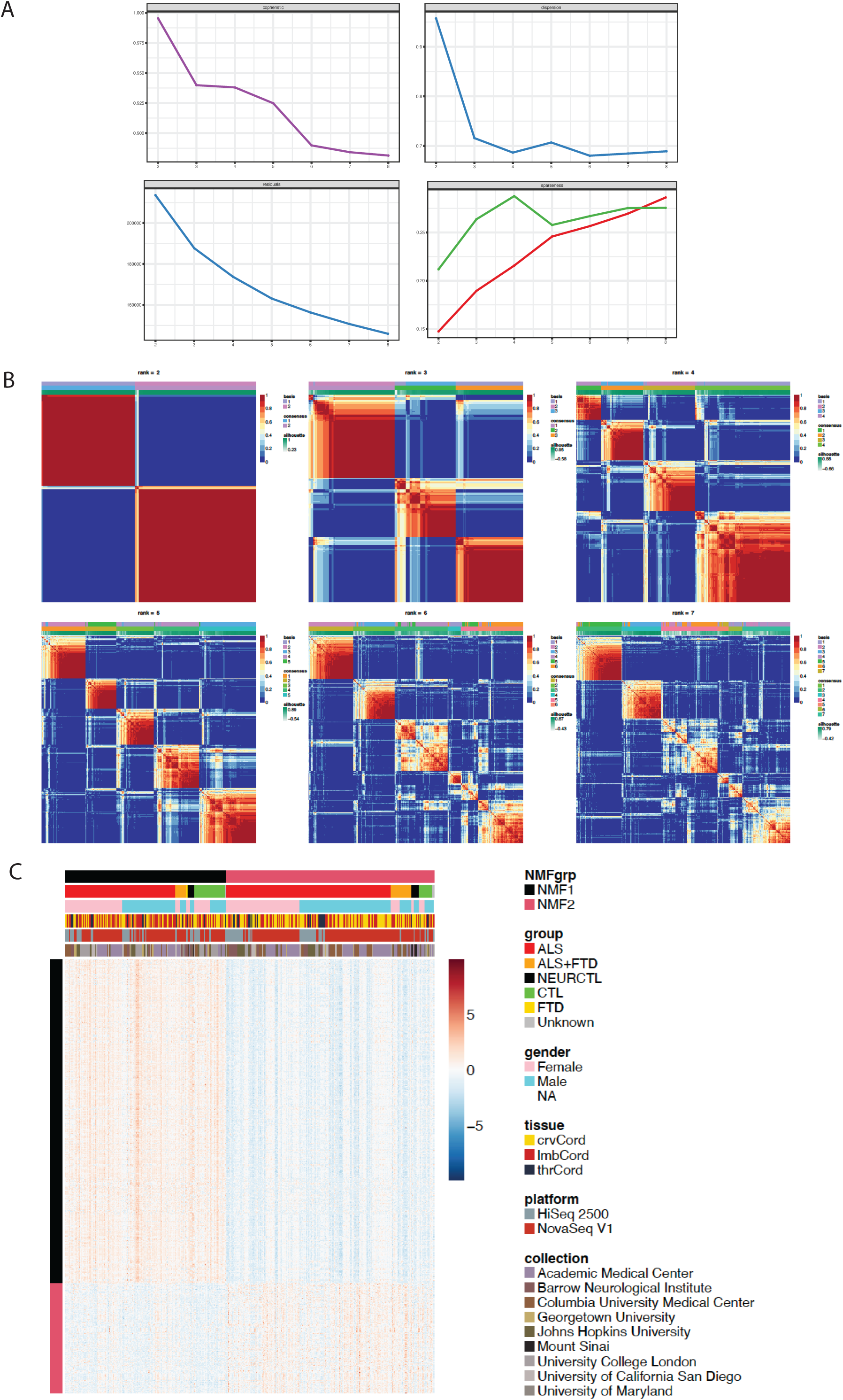
Non-negative matrix factorization of spinal cord samples. A) Cophenetic (top left), dispersion (top right), residual (bottom left) and sparseness (bottom right) for each rank value tested. B) Consensus map of NMF clusters at rank 2 to 7. Rank of 2 was subsequently chosen due to high cophenetic and low sparseness. C) Expression heatmap of NMF marker genes in frontal/motor cortex samples. NMF assignment does not show obvious correlation with gender, tissue type, sequencing platform or collection site.

**Figure S4:**
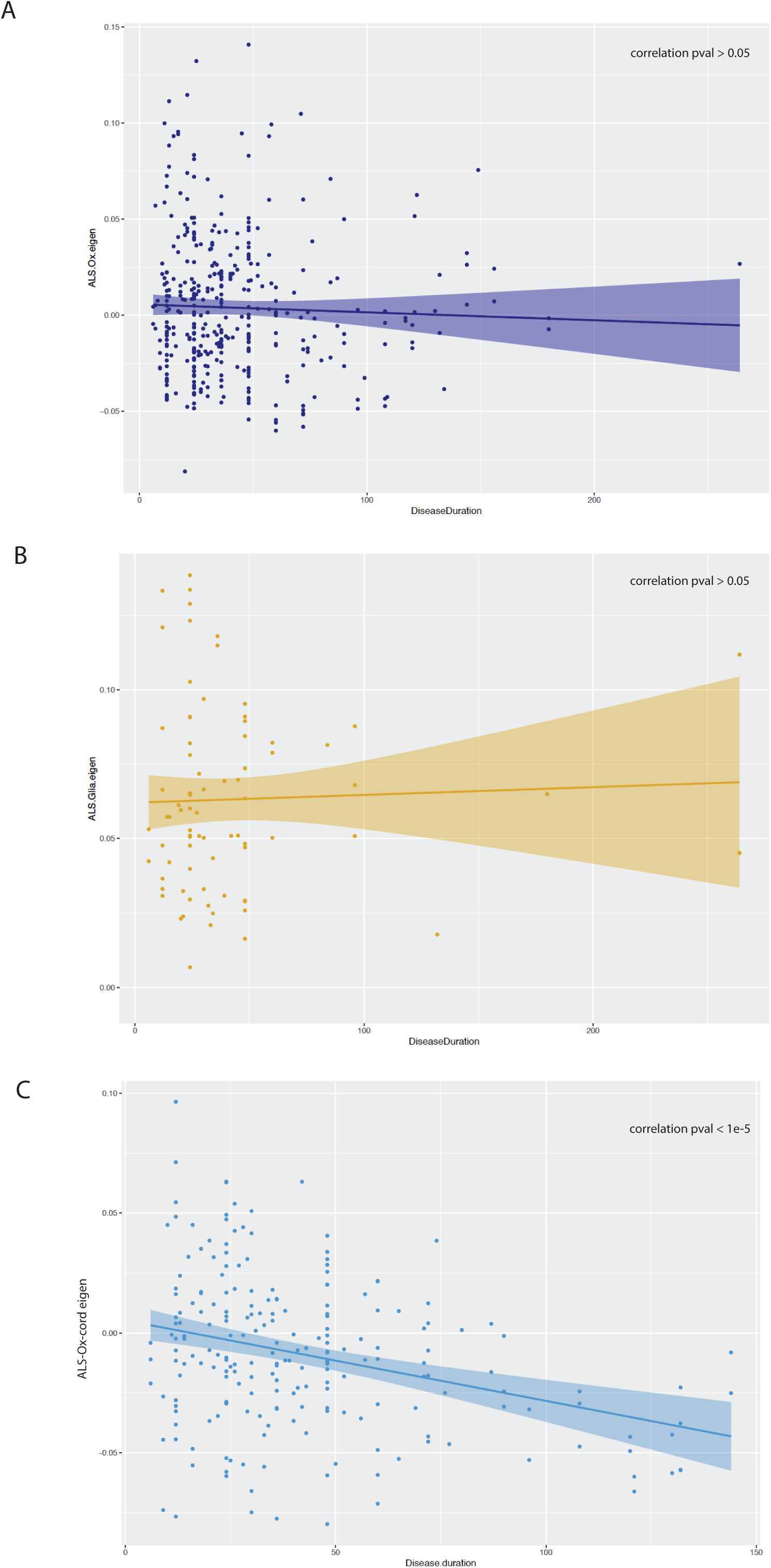
Correlation of ALS subtype features with disease duration. Scatterplot of eigengene value representing ALS subtype features for ALS-Ox (A), ALS-Glia (B) and ALS-Ox-cord (C) with disease duration. No significant correlation between the two variables were observed.

**Figure S5:**
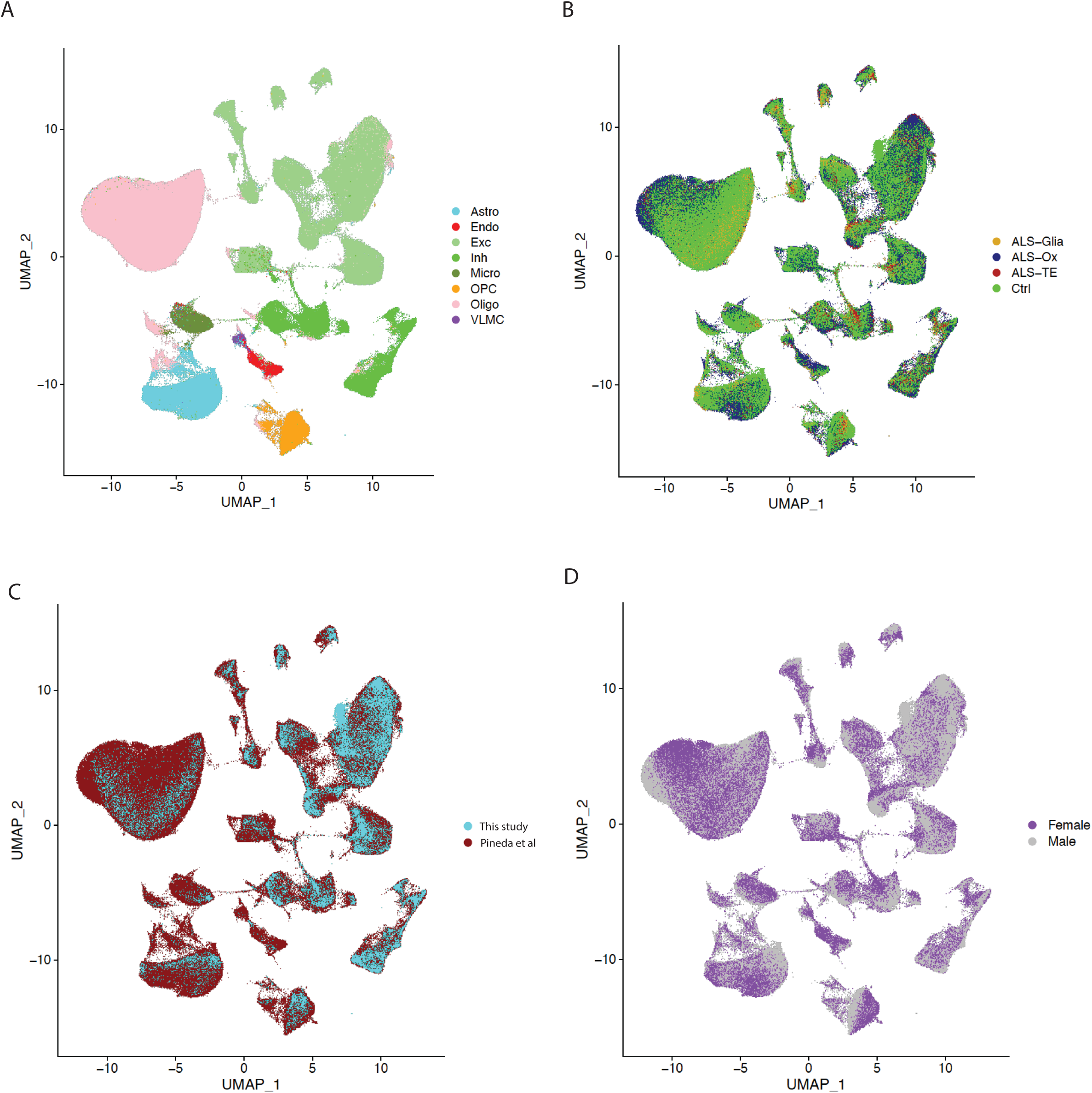
UMAP projection of integrated single-nuclei datasets. Clusters show strong association with cell type (A), but not ALS molecular subtype (B), dataset source (C), nor sex (D)

**Figure S6:**
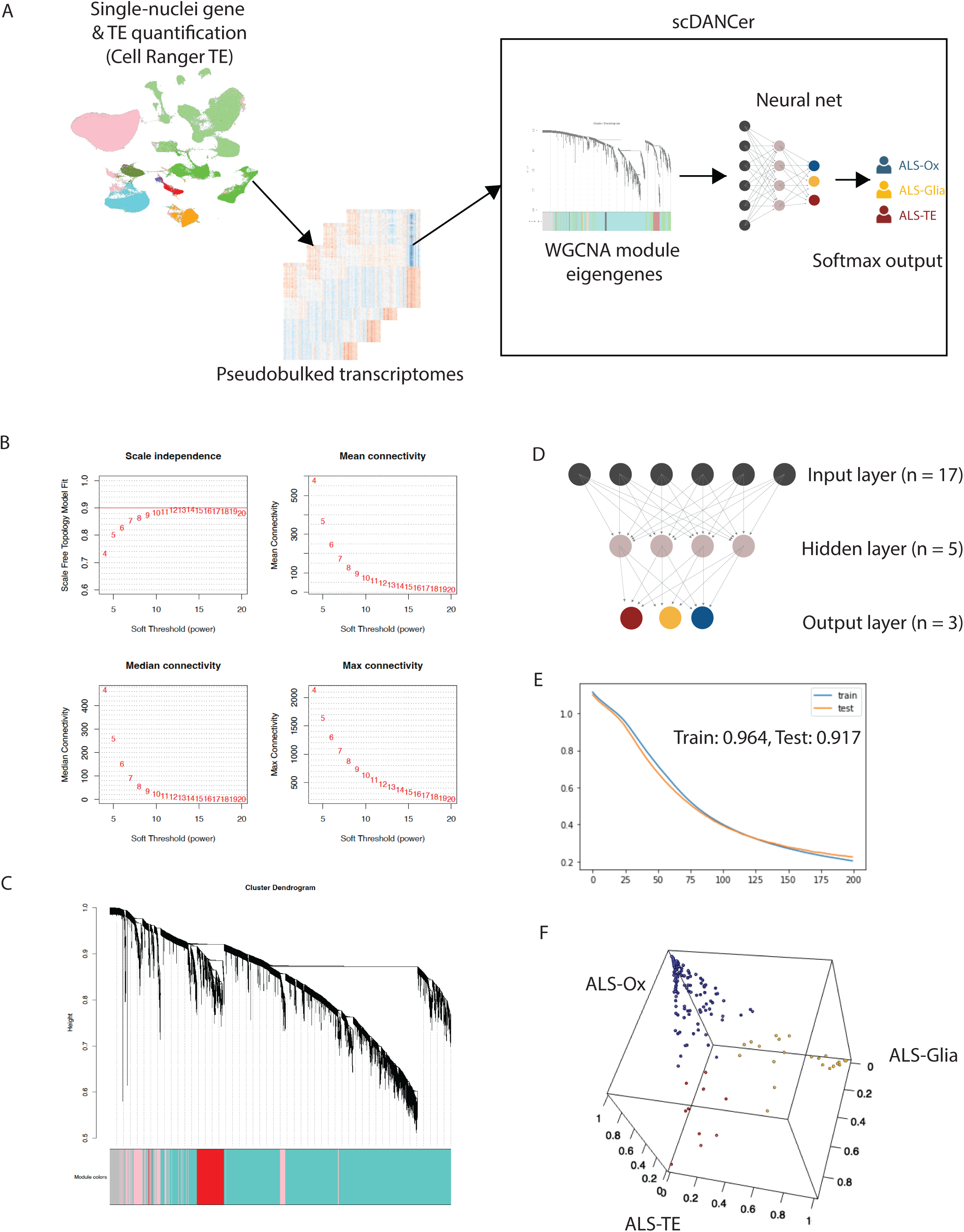
single-cell Deep ALS Neural-Net Classifier (scDANCer). A) Workflow to utilize scDANCer. Gene and TE quantification is done using Cell Ranger TE, followed by pseudobulking of cellular transcriptome for each sample. After variance stabilized transformation, module eigengene is calculated, and the fed into the neural net for ALS subtype classification. B) Determining the best soft threshold/power parameter for WGCNA. The power parameter of 9 was chosen due to a scale-free model correlation close to 0.9 while maintaining connectivity within the network. C) Hierarchical clustering of genes and their assignment to WGCNA modules. D) Layout of the neural net, with 17 nodes in the input layer, 5 nodes in the hidden layer and 3 softmax nodes in the output layer. E) Plot of categorical cross entropy loss in the training and test sets after each epoch of training. The final accuracy is 97.2% for the training set and 91.7% for the test set. F) 3D plot of the classifier confidence from the softmax output, with the dots labeled by their final ALS subtype classification.

**Table S1: Metadata for frontal/motor cortex samples**. A) Summary of metadata features. B) Metadata and ALS subtype for frontal/motor cortex samples in this study. C) Clinical metadata and ALS subtype for patients with frontal/motor cortex samples in this study.

**Table S2: Association of ALS subtype with technical and clinical features**. A) Comparison of pH and RIN between subtypes and controls in frontal/motor cortex samples. B) Association of ALS subtype with various clinical features

**Table S3: Metadata for spinal cord samples** A) Summary of metadata features. B) Metadata and molecular subtype for spinal cord samples in this study. C) Clinical metadata and molecular subtype for patients with spinal cord samples in this study. D) Association of subtype with sample pH & RIN

**Table S4: Differential analysis and GSEA of ALS molecular subtypes**. Tables contain differentially expressed features between ALS-Glia (A), ALS-Ox (C), ALS-TE (E), ALS-Inflam-cord (G) or ALS-Ox-cord (I) vs control, or significantly enriched/depleted gene sets from GSEA between ALS-Glia (B), ALS-Ox (D), ALS-TE (F), ALS-Inflam-cord (H) or ALS-Ox-cord (J) vs control.

**Table S5: Clinical metadata and molecular subtype of patients with frontal/motor cortex and spinal cord samples in this study**

**Table S6: Differential splicing results of ALS molecular subtypes**. Tables contain differentially spliced junctions between ALS-Glia (A), ALS-Ox (B), ALS-TE (C), ALS-Inflam-cord (D) or ALS-Ox-cord (E) vs control.

**Table S7: Evaluating subtype calls between DANCer and scDANCER.** A) Comparison of subtypes called by DANCer and scDANCER from bulk and down-sampled NYGC RNAseq dataset from Tam et al. 2019. B) Comparison of subtypes called by DANCer of non-NYGC bulk RNAseq dataset from Tam et al. with corresponding snRNA-seq dataset generated in this study. C) Comparison of subtypes called by DANCer between two replicates from the Pineda et al. 2024 dataset.

**Table S8: Differential analysis and GSEA of L5ET cells from scDANCer classified samples.** Tables contain differentially expressed features from ALS-Glia (A), ALS-Ox (C) or ALS-TE (E) L5ET cells vs control, or enriched/depleted gene sets from ALS-Glia (B), ALS-Ox (D) or ALS-TE (F) L5ET cells vs control. G) GSEA of top 200 up/down-regulated genes from TDP-43 depleted neuronal nuclei (Liu et al. 2018). H) Differentially expressed features common to all three subtypes vs control

**Table S9: Differential analysis and GSEA of astrocytes from scDANCer classified samples.** Tables contain differentially expressed features from ALS-Glia (A), ALS-Ox (C) or ALS-TE (E) astrocytes vs control, or GSEA enriched/depleted gene sets from ALS-Glia (B), ALS-Ox (D) or ALS-TE (F) astrocytes vs control. G) GSEA of top 200 up/down-regulated genes from TDP-43 depleted neuronal nuclei (Liu et al. 2018). H) Differentially expressed features common to all three subtypes vs control

**Table S10: Differential analysis and GSEA of microglia from scDANCer classified samples.** Tables contain differentially expressed features from ALS-Glia (A), ALS-Ox (C) or ALS-TE (E) microglia vs control, or GSEA enriched/depleted gene sets from ALS-Glia (B), ALS-Ox (D) or ALS-TE (F) microglia vs control. G) GSEA of top 200 up/down-regulated genes from TDP-43 depleted neuronal nuclei (Liu et al. 2018). H) Differentially expressed features common to all three subtypes vs control

